# Astrocytic sEH–EET signaling regulates microglial lysosomal function and suppresses the STING–Type I interferon pathway in Alzheimer’s disease

**DOI:** 10.64898/2026.10.16.759679

**Authors:** Baijayanti Ghosh, Shuo Wang, Chuangye Qi, Anamitra Ghosh, Michele Comerota, Chetan Singh Rajpurohit, Darryl C. Zeldin, Hui Zheng

**Affiliations:** Department of Molecular and Human Genetics, Baylor College of Medicine, Houston, Texas, USA; National Institute of Environmental Health Sciences, National Institutes of Health, Research Triangle Park, North Carolina, USA; Department of Neuroscience, Baylor College of Medicine, Houston, Texas, USA

**Keywords:** Alzheimer’s disease, β-amyloid, microglia, soluble epoxide hydrolase, STING

## Abstract

**Background:** The development of β-amyloid (Aβ) pathology in Alzheimer’s disease (AD) is accompanied by profound changes in astrocytes and microglia and associated neuroinflammation. Soluble epoxide hydrolase (sEH), which degrades anti-inflammatory epoxyeicosatrienoic acids (EETs), is elevated in the AD brain, yet how EETs restrain inflammation remains poorly understood.

**Methods:** We performed single-nucleus RNA sequencing (snRNA-seq) on brains from wild-type, sEH knockout, 5xFAD, and 5xFAD; sEH knockout mice, and used primary mouse microglia and human induced pluripotent stem cell (iPSC)-derived microglia to define the underlying mechanism. To test cell-type-specific contributions, we generated astrocyte- and endothelial-specific she knockout mice and crossed them with 5xFAD mice.

**Results:** Genetic disruption of sEH in 5xFAD mice markedly reduced the abundance of a type I interferon (IFN)-responsive microglial subcluster and lowered phospho-STING (pSTING) levels. In primary microglia, supplementation with 11,12-EET attenuated STING activation and IFN target gene expression; blocking lysosomal acidification abolished this effect and caused pSTING to accumulate, identifying lysosomal degradation as a critical route for terminating STING signaling. 11,12-EET also enhanced microglial lysosomal acidification and Aβ phagocytosis. Astrocytic, but not endothelial, disruption of sEH recapitulated the reductions in Aβ pathology and microglial STING activation seen in global sEH-deficient mice, identifying astrocytes as the major cellular source of EETs regulating this pathway.

**Conclusion:** We define an astrocyte–microglia axis in which astrocytic sEH limits EET availability and thereby sustains microglial STING signaling. By promoting lysosomal clearance of pSTING, EETs suppress type I IFN signaling and enhance microglial Aβ clearance, highlighting sEH inhibition as a therapeutic strategy targeting both neuroinflammation and lysosomal dysfunction in AD.

## Introduction

Alzheimer’s disease (AD) is a progressive neurodegenerative disorder defined by extracellular amyloid-β (Aβ) plaques, intracellular tau tangles, synaptic failure, and neuronal loss. Although Aβ accumulation is a central pathological hallmark, growing evidence indicates that glial cells and innate immune responses critically shape disease trajectory [1–4]. Recent antibody-based trials targeting Aβ yielded only modest clinical benefit, underscoring the need for complementary strategies that address immune and inflammatory mechanisms operating within the diseased brain [5–7].

Microglia display marked phenotypic heterogeneity in AD, transitioning from homeostatic states toward transcriptionally distinct disease-associated microglia (DAM) and Type I interferon (IFN)–responsive states enriched around plaques [8–10]. Single-cell and single-nucleus RNA-sequencing studies have linked these states to aberrant lipid metabolism and lysosomal activity [8–13]. IFN-responsive microglia in particular contribute to neuritic dystrophy, synapse loss, and cognitive impairment downstream of amyloid deposition, implicating their role in disease progression [14, 15].

The cGAS–STING axis has emerged as a key regulator of neuroinflammatory tone in aging and neurodegeneration [16–18]. Aberrant STING activation in microglia promotes sustained Type I IFN production and neurotoxicity [19]. Termination of STING signaling requires its trafficking of STING to lysosomes, and disrupting lysosomal acidification or degradative capacity prolongs STING activation and inflammatory signaling [20–24]. These findings position the lysosome as a critical checkpoint of STING signaling.

Soluble epoxide hydrolase (sEH, encoded by the *Ephx2* gene) metabolizes epoxy-fatty acids, including epoxyeicosatrienoic acids (EETs), which exert anti-inflammatory and neuroprotective actions [25–28]. sEH is broadly expressed across peripheral tissues, including liver, kidney, and vasculature, where the EET-sEH axis regulates endothelial function, vascular homeostasis, and inflammatory responses [29–35]. Within the central nervous system, sEH is expressed in multiple cell types and is particularly enriched in astrocytes, in which its expression is increased in AD brain and in experimental amyloidosis [36]. Genetic or pharmacologic inhibition of sEH elevates endogenous EET levels, suppresses neuroinflammation, attenuates amyloid pathology, and improves synaptic and cognitive outcomes in AD mouse models [36–38]. More recently, Wu et al. showed that an age-associated increase in hepatic sEH lowers circulating 11,12-EET levels and influences cerebral Aβ burden, tau pathology, and cognitive impairment, revealing a previously unrecognized liver-brain axis in AD pathogenesis [39]. Together, these studies establish sEH as a regulator of both peripheral and central mechanisms relevant to neurodegeneration.

Despite accumulating evidence for therapeutic benefits from sEH inhibition, the mechanisms linking sEH-EET signaling to glial immune state transitions remain incompletely understood. Herein, we investigated the sEH-EET axis in immune regulation and test the cell-type-specific contribution of sEH in 5XFAD mice. We find that sEH modulates glial immune states, lysosomal competence, and STING-IFN signaling, and we identify astrocytic, but not endothelial sEH as a key upstream regulator of these processes.

## Materials and methods

### Study design

The objective of this study was to determine how the soluble epoxide hydrolase (sEH)-epoxyeicosatrienoic acid (EET) signaling pathway regulates glial immune states, lysosomal competence, and amyloid pathology in Alzheimer’s disease. To address this question, we combined germline, astrocyte-specific, and endothelial-specific disruption of *Ephx2* with single-nucleus RNA sequencing, immunohistochemistry, biochemical analyses, behavioral testing, primary glial cultures, and human induced pluripotent stem cell (iPSC)-derived microglia-like cells (iMGLs). Pharmacological and biochemical approaches were used to investigate the effects of EET signaling on STING activation, lysosomal function, and Aβ processing.

Animals, cell cultures, imaging experiments, and quantitative analyses were randomized when appropriate and performed with investigators blinded to genotype and treatment group. Sample sizes for animal studies were determined based on previous studies and experience with the 5XFAD model [36]. The exact number of animals used in each experiment is reported in the corresponding figure legends. Behavioral analyses are presented in the Supplementary Information. All in vitro experiments were performed using at least three biological replicates and were independently repeated at least twice. Additional experimental details are provided in the sections below and in the corresponding figure legends.

### Mice

The 5XFAD mice (referred to as 5X in this study) were obtained from The Jackson Laboratory (JAX Strain# 034840). Global sEH knockout (*Ephx2^-/-^,* referred to as sEH KO in this study) mice were maintained on a C57BL/6J background and crossed with 5XFAD mice to generate 5X sEH KO animals. Conditional *Ephx2* floxed mice were generated as previously described by Bettaieb et al. and maintained on a C57BL/6J background [40]. *Ephx2^fl/fl^* mice were crossed with GFAP-Cre mice (JAX strain# 24098) to generate astrocyte-specific knockout mice (referred to as sEH AKO in this study) [41] and with Cdh5-Cre (JAX strain# 006137) mice to generate endothelial-specific knockout mice (referred to as sEH EKO in this study) [42]. These lines were subsequently crossed with 5XFAD mice to generate astrocyte-specific sEH knockout (5X sEH AKO) and endothelial-specific sEH knockout (5X sEH EKO) experimental cohorts. Unless otherwise indicated, all experiments were performed in 6-month-old mice. Both male and female animals were used and were balanced across experimental groups. Mice were housed in a pathogen-free facility with ad libitum access to food and water under a 12-h light/dark cycle. All experimental procedures were approved by the Baylor College of Medicine Institutional Animal Care and Use Committee (IACUC) and conducted in accordance with NIH guidelines.

### Single nucleus RNA-sequencing and data analysis

Single-nucleus RNA sequencing (snRNA-seq) was performed as previously described [43]. Briefly, 6-month-old WT, sEH KO, 5XFAD, and 5XFAD sEH KO mice (2 males and 2 females per group) were transcardially perfused with ice-cold saline. Hippocampi were rapidly dissected, dissociated, and nuclei were isolated. Nuclei were stained with Hoechst-33342 and sorted using a SONY SH800 fluorescence-activated cell sorter. For each 10x Genomics run, 100,000–400,000 nuclei were collected, and 16,000 nuclei per channel were loaded into the 10x Chromium Controller.

snRNA-seq libraries were generated using the 10x Genomics Chromium Single Cell 3′ v3.1 kit, following the manufacturer’s protocol. All PCR steps were performed on a Bio-Rad C1000 Touch Thermal Cycler equipped with a 96-Deep Well Reaction Module. Thirteen PCR cycles were used for cDNA amplification and sixteen cycles for sample index amplification. As recommended by 10x Genomics, 1:10 dilutions of amplified cDNA and final libraries were quality-controlled using a Bioanalyzer. Male and female samples were pooled and sequenced using 100-cycle run kits in a single-index configuration. Gene expression libraries were sequenced using paired-end 150-bp reads on an Illumina NovaSeq platform to achieve ∼60–70% sequencing saturation.

Raw sequencing data were demultiplexed using bcl2fastq, and gene expression libraries were processed using Cell Ranger (v7.1.0) with the mm10 reference genome. Ambient RNA contamination was estimated and removed using CellBender [44]. Doublets were identified using Scrublet [45] applied to the filtered feature-barcode matrices generated by CellBender, and nuclei with doublet scores > 0.25 were excluded. Additional filtering was performed using Scanpy (v1.9.5) [46] retaining nuclei with < 5% mitochondrial UMI content and > 200 detected genes.

Batch correction and low-dimensional embeddings were generated using single-cell variational inference (scVI) [47]. Major cell types and subclusters were annotated based on established marker genes. Spliced and unspliced transcript counts were obtained using velocyto (run10x) and were normalized and smoothed over 15 nearest neighbors using the filter_and_normalize and moments functions in scVelo [48, 49]. RNA velocity, transition matrix estimation, and streamline visualization were performed using the velocity and velocity_embedding_stream functions with dynamical modeling. Normalized spliced and unspliced counts were used for pseudotime trajectory inference and visualization in UMAP space. Differential gene expression analysis between experimental groups and microglial subclusters was performed using the Wilcoxon rank-sum test implemented in the FindMarkers function [50]. Genes were considered differentially expressed if they met the following criteria: false discovery rate (FDR) < 0.05, average log2 fold change > 0.1, and expression in > 10% of cells in at least one group.

Single-cell regulatory network inference and clustering (SCENIC) analysis was performed using pySCENIC (v0.12.1) to infer transcription factor (TF)-target gene regulatory networks in microglia from snRNA-seq data. The analysis was conducted using the raw count matrix of microglial nuclei from all samples, together with the Mus musculus mm10 transcription start site (TSS)-centered ±10 kb regulatory databases available from the pySCENIC repository [51]. The workflow consisted of three sequential steps. First, GRNBoost was used to infer TF-target gene co-expression modules and estimate the regulatory importance of candidate target genes. Second, RcisTarget was applied to identify enriched TF-binding motifs and refine the co-expression modules into regulons comprising TFs and their putative direct targets. Third, AUCell was used to quantify the activity of each regulon in individual cells based on the expression of its target genes. For visualization, mean regulon AUC scores were calculated for the relevant microglial populations, and TF-target regulatory networks (in IFN microglia) were drawn using NetworkX with its networkx.drawing.nx_agraph interface.

### Primary microglia cultures and treatment

Primary microglia cultures were prepared from postnatal day 1-3 mouse pups as described previously [52]. Cortices and hippocampi were dissected, minced, and enzymatically digested in 2.5% trypsin at 37 °C for 15 min. Tissue was then centrifuged, mechanically triturated, and resuspended in DMEM supplemented with 10% fetal bovine serum (FBS). Cells were plated onto poly-D-lysine (PDL)-coated T-75 flasks to establish mixed glial cultures. Cultures were maintained for two weeks, with medium replenished every other day.

Microglia were isolated from mixed glial cultures using CD11b magnetic microbead selection according to the manufacturer’s instructions (Miltenyi Biotec). Enriched microglia were plated onto PDL-coated 24-well plates in DMEM containing 10% FBS and 1% penicillin/streptomycin, supplemented with 10 ng/mL macrophage colony-stimulating factor (M-CSF). Cells were allowed to recover for 24-48 h prior to experimental treatments.

Microglia were treated with vehicle (DMSO) or 11,12-EET (10 µM) for 18 h. For Aβ42 phagocytosis assays, pHrodo™ Red-conjugated Aβ42 (500 ng/mL) was prepared following the manufacturer’s instructions (FUJIFILM Cellular Dynamics). Lyophilized peptide was dissolved in DMSO and added to the cultures for an additional 1 h. The peptide-containing medium was then removed and replaced with fresh medium, and cells were washed with PBS. One set of cultures was fixed after 1 hour in 4% paraformaldehyde to assess Aβ uptake, while the second set was incubated in fresh medium for an additional 24 h to evaluate Aβ clearance (Fig. S4). Images were acquired using a Leica STELLARIS confocal microscope with a 40× objective.

To assess the effects of EET on STING activation, microglia were treated with 11,12-EET (10 μM; Cayman Chemical) 2 hours prior to adding the murine STING agonist DMXAA (10 μg/mL; Invivogen, treated for 4h). For lysosomal inhibition studies, cells were co-treated with bafilomycin A1 (BAF; 50 nM; Sigma-Aldrich) and DMXAA. Cells were harvested after 4 h of DMXAA, with or without BAF treatment, for immunofluorescence, immunoblot analyses and quantitative PCR analyses of interferon-stimulated genes (ISGs).

### iMicroglia culture and treatment

Human induced pluripotent stem cell (iPSC)-derived microglia-like cells (iMGLs) were generated using a modified version of previously published protocols by Abud et al. and McQuade et al [53, 54]. Human iPSCs of early passage (male, GM23815, 196 iPSCs clone-13) were acquired from the Baylor College of Medicine (BCM) Stem Cell Core and cultured in Matrigel-coated dishes (Millipore Cat# CLS354234) in mTeSR Plus medium (Stem Cell Technologies Cat#100-0276). iPSCs were differentiated into hematopoietic progenitor cells (HPCs) using the STEMdiff™ Hematopoietic Kit (Stemcell Technologies) and CD43⁺ HPCs were isolated by fluorescence-activated cell sorting. HPCs were subsequently differentiated into microglia-like cells in serum-free DMEM/F12-based medium supplemented with IL-34 (100 ng/mL), TGF-β1 (50 ng/mL), and M-CSF (25 ng/mL). During the final maturation phase, cells were cultured in the presence of CD200 (100 ng/mL) and CX3CL1 (100 ng/mL) to promote acquisition of a mature microglial phenotype. Differentiation cultures were maintained for approximately 40 days, after which iMGLs were used for experiments. The differentiation protocol was adapted from previously published methods and optimized in-house as described by Abud et al. and McQuade et al [53, 54]. For STING activation studies, iMGLs were treated with 11,12-EET (10 μM), bafilomycin A1 (100 nM), the human-selective STING agonist diABZI (1 μM, Selleck Chemicals); while EET was added 2h prior to diABZI treatment, Bafilomycin was added along with diABZI for all experiments. Cells were fixed after 4h of treatment of dIABZI for immunofluorescence analyses.

### Western blotting

Cells or brain tissues were lysed in modified radioimmunoprecipitation assay (RIPA) buffer supplemented with protease and phosphatase inhibitor cocktails. Cell lysates were vortexed briefly following resuspension, whereas mouse brain tissues were homogenized, vortexed, and centrifuged at 14,000 × g for 45 min at 4°C, as previously described [36]. Protein concentrations were determined using a bicinchoninic acid (BCA) assay (Thermo Fisher Scientific). Equal amounts of protein were resolved on 7.5–15% SDS–polyacrylamide gels (Bio-Rad) and transferred onto nitrocellulose membranes. Membranes were blocked with Tris-buffered saline (TBS) containing 5% bovine serum albumin (BSA) and subsequently incubated with primary antibodies followed by appropriate secondary antibodies. Immunoreactive bands were visualized and quantified using Bio-Rad Image Lab software, and protein levels were normalized to loading controls. Primary antibodies used for western blotting are listed in Supplemental Table 1.

### RNA extraction and expression analysis

Total RNA was extracted from cultured cells or brain tissues using the RNeasy Mini Kit (Qiagen, 74106). Reverse transcription was performed using iScript Reverse Transcription Supermix (Bio-Rad, 1708840) to generate cDNA. Quantitative PCR was carried out using SYBR Green Master Mix (Bio-Rad) on a CFX384 Touch Real-Time PCR Detection System (Bio-Rad). Relative gene expression was calculated using the ΔΔCt method and normalized to the housekeeping gene *Gapdh*. Primers used have been enlisted in Supplemental Table 2.

### Immunofluorescence staining

For tissue staining, mice were deeply anesthetized and transcardially perfused with PBS, followed by fixation in 4% paraformaldehyde (PFA) overnight at 4 °C. Brains were cryoprotected in 30% sucrose in PBS, and coronal sections (30 µm) were cut using a Leica sliding microtome. Free-floating sections were blocked in 10% normal donkey serum in PBS containing 0.1% Triton X-100 for 1 h at room temperature, then incubated with primary antibodies diluted in blocking solution overnight at 4°C. Sections were washed and incubated with fluorescent secondary antibodies and DAPI in blocking buffer for 1 h at room temperature, followed by mounting with Fluoromount-G.

For cultured cells, primary cultures grown on coverslips were fixed with 4% PFA for 20 min, permeabilized with 0.1% Triton X-100 in PBS for 5 min, and blocked in 4% normal donkey serum containing 0.01% Triton X-100 for 1 h at room temperature. Coverslips were incubated with primary antibodies diluted in blocking buffer overnight at 4°C, washed with PBS, and incubated with fluorescent secondary antibodies and DAPI for 1–1.5 h at room temperature. After final washes, coverslips were mounted onto glass slides using ProLong™ Gold antifade reagent. Primary antibodies used for immunofluorescence are enlisted in Supplemental Table 1.

### Image acquisition and analysis

All image acquisition and quantification were performed with investigators blinded to genotype and treatment condition. Images within a given experiment were acquired using identical laser power, detector gain, offset, and exposure settings. Coronal brain sections (30 μm) and cultured cells were imaged using a Leica STELLARIS confocal microscope. For tissue analyses, confocal z-stacks spanning the entire thickness of each section were acquired at 1 μm intervals. Images were processed and analyzed using Fiji/ImageJ (NIH) and Imaris software (Bitplane, Oxford Instruments).

### Mouse brain

For histological analyses, at least three anatomically matched hippocampal sections per animal were quantified and averaged to generate one biological replicate. Fluorescence-positive area (%) and mean fluorescence intensity were quantified in Fiji/ImageJ following background subtraction using identical thresholding parameters across all groups within an experiment.

Amyloid plaque burden was quantified in Imaris by generating three-dimensional plaque surfaces from Methoxy-X04-positive signals using the Surface module. Plaque number and plaque size were then calculated from the rendered surfaces.

For analyses of plaque-associated microglia, Methoxy-X04-positive plaques and IBA1-positive microglia were rendered as three-dimensional surfaces and spots respectively in Imaris. Microglial spots located nearest to amyloid plaque surfaces (within 20μm) were identified using the Shortest Distance filter and selected for subsequent analyses.

To assess pSTING and pSTAT1 localization within microglia, regions of interest (ROIs) were generated for IBA1-positive microglia and pSTING- or pSTAT1-positive signal. Colocalization analysis was performed using the Imaris Coloc module. Microglial STING and STAT1 activation were quantified as the percentage of ROI material colocalized (% ROI Material Colocalized) between IBA1 and pSTING or between IBA1 and pSTAT1, respectively. Identical thresholding and colocalization parameters were applied to all images within a given experiment.

### Cell culture

For primary microglia and iPSC-derived microglia-like cells (iMGLs), fluorescence intensity was quantified on a per-cell basis using Fiji/ImageJ after background subtraction. Multiple imaging fields were analyzed for each biological replicate and averaged prior to statistical analysis.

For AF488 bead uptake, pHrodo-Aβ, DQ-BSA, and Aβ42 degradation assays, fluorescence intensity was quantified using Fiji/ImageJ. For pHrodo-Aβ assays, fluorescence intensity measured 1 h after addition of pHrodo-Aβ was used as an indicator of Aβ uptake and lysosomal delivery, whereas fluorescence intensity measured 24 h later was used as an assessment of intracellular Aβ retention and degradation. Data from multiple fields were averaged to generate a single biological replicate for statistical analysis.

## Statistical analysis

All statistical analyses were performed using GraphPad Prism (v10.4.1). Data are presented as mean ± SEM, with statistical significance defined as \**p* < 0.05, \*\**p* < 0.01, \*\*\**p* < 0.001, and \*\*\*\**p* < 0.0001. For pairwise comparisons, Welch’s *t*-test was applied. Comparisons involving multiple groups were analyzed using ordinary one-way ANOVA or Welch’s ANOVA followed by Tukey’s post hoc test or Games-Howell test respectively. All samples and animals were included in the analyses unless otherwise stated.

## Results

### sEH disruption remodels microglial states

Previous studies demonstrated that inhibition of sEH confers anti-inflammatory effects in AD mouse models [36, 37]. To define associated microglial transcriptional changes, we crossed the global *Ephx2* knockout mice (herein referred to as sEH KO) with 5XFAD mice. Consistent with pharmacological inhibition, genetic disruption of the *Ephx2* gene resulted in reduced Aβ plaques and reactive gliosis, diminished inflammatory cytokines and improved behavior (Supplementary Fig. S1). We performed single-nucleus RNA sequencing (snRNA-seq) on hippocampal tissue from WT, sEH KO, 5XFAD, and 5XFAD sEH KO mice. Following quality control and integration, unsupervised clustering of a total 76,119 nuclei identified major brain cell populations, including excitatory neurons, inhibitory neurons, astrocytes, oligodendrocytes, oligodendrocyte precursor cells, microglia, and vascular cells (Fig. 1A). Visualization by UMAP demonstrated clear segregation of these populations. Batch correction effectively integrated samples across genotypes and sex, as indicated by the even distribution of nuclei when categorized by genotype or sex (Fig. S2, A & B). Visualization of individual genotypes confirmed similar clustering patterns across WT, sEH KO, 5XFAD, and 5XFAD sEH KO samples (Fig. S2, C-F). Quantification of major cell-type proportions revealed no substantial differences between genotypes (Fig. 1B), indicating that germline sEH disruption does not grossly alter cellular composition in the hippocampus.

**Figure 1.**
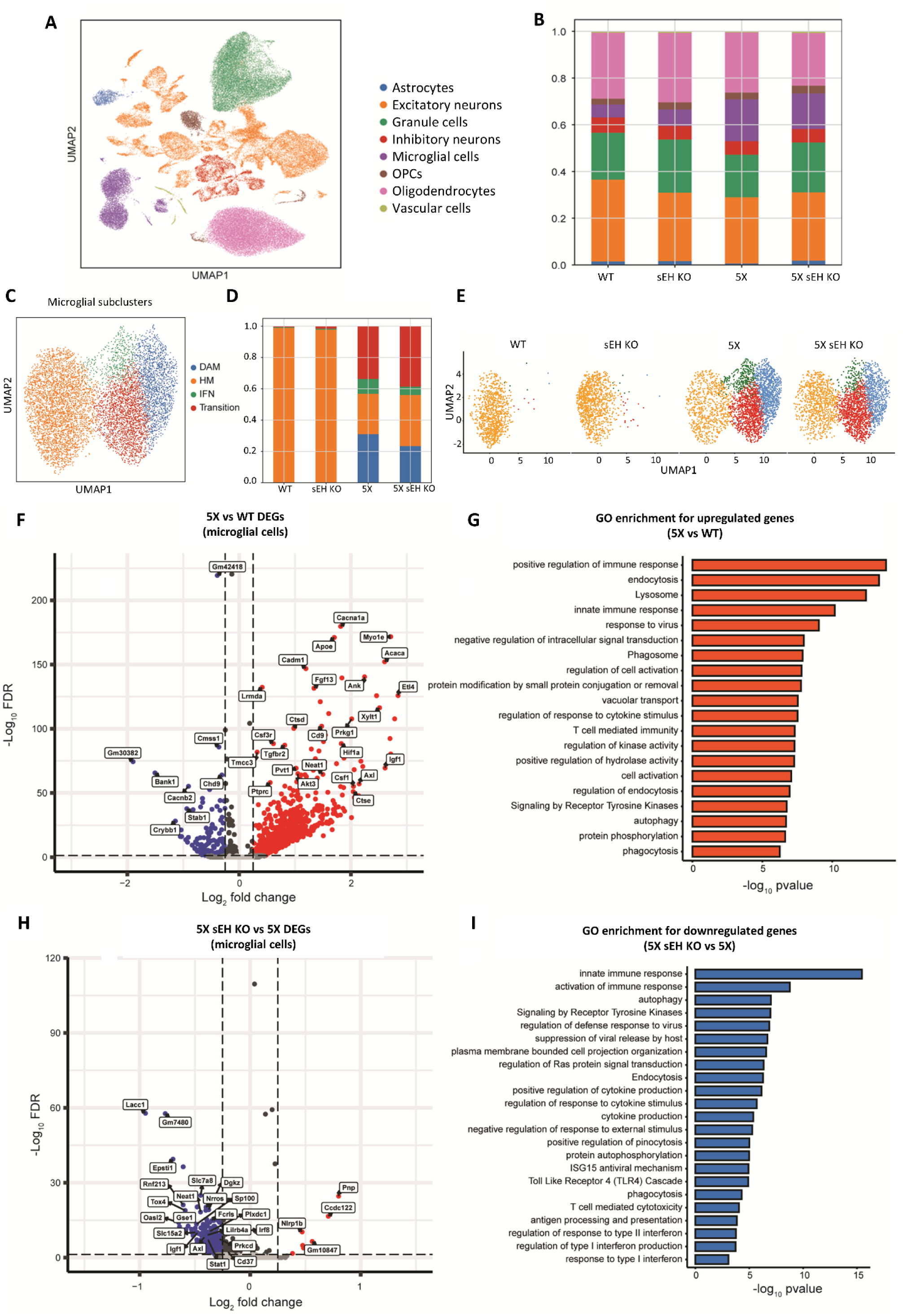
sEH deletion remodels microglial transcriptional states and suppresses interferon-associated gene programs. (A) UMAP embedding of snRNA-seq data showing 76,119 cells from the mice hippocampus of four genotypes (WT, sEH KO, 5XFAD (5X), and 5XFAD sEH KO (5X sEH KO) at 6-month-old, annotated by cluster. (B) Bar chart representation of the proportion of different cell types identified in the snRNA-seq across different genotypes. (C) UMAP embedding of snRNA-seq data showing 7,839 microglial cells from the mice hippocampus of four genotypes at 6-month-old, annotated by microglial subtypes. HM: homeostatic microglia; IFN: interferon microglia; DAM: Disease-associated microglia. (D) Bar chart representation showing the proportion of microglial subtypes across different genotypes. (E) UMAP embedding of snRNA-seq data for microglial subtypes across different genotypes. (F) Volcano plot showing the differentially expressed genes for microglial cells in 5xFAD relative to WT mice. Upregulated genes are highlighted in red color. Downregulated genes are highlighted in blue color. (G) GO pathway enrichment for the upregulated genes for microglial cells in 5xFAD relative to WT mice. The enriched pathways are highlighted in orange color. (H) Volcano plot showing DEGs in microglia comparing 5XFAD sEH KO versus 5XFAD, with representative genes indicated. (I) GO enrichment analysis of genes downregulated in 5XFAD sEH KO relative to 5XFAD, showing top enriched biological processes ranked by −log₁₀ p value.

We next focused on microglial populations, given their close association with plaque pathology. Reclustering of a total 7,839 microglial nuclei identified multiple transcriptionally distinct microglial subclusters (Fig. 1C, Fig. S2G), including homeostatic (HM), transitional, disease-associated (DAM), and interferon-responsive (IFN). Comparison of subcluster distributions across genotypes revealed an expansion of DAM and IFN microglial populations in 5XFAD mice, which was reduced in 5XFAD sEH KO mice (Fig. 2D). Consistently, UMAP visualization colored by genotype demonstrated an altered microglial state distribution in 5XFAD mice that was partially normalized by sEH loss (Fig. 1E).

**Figure 2.**
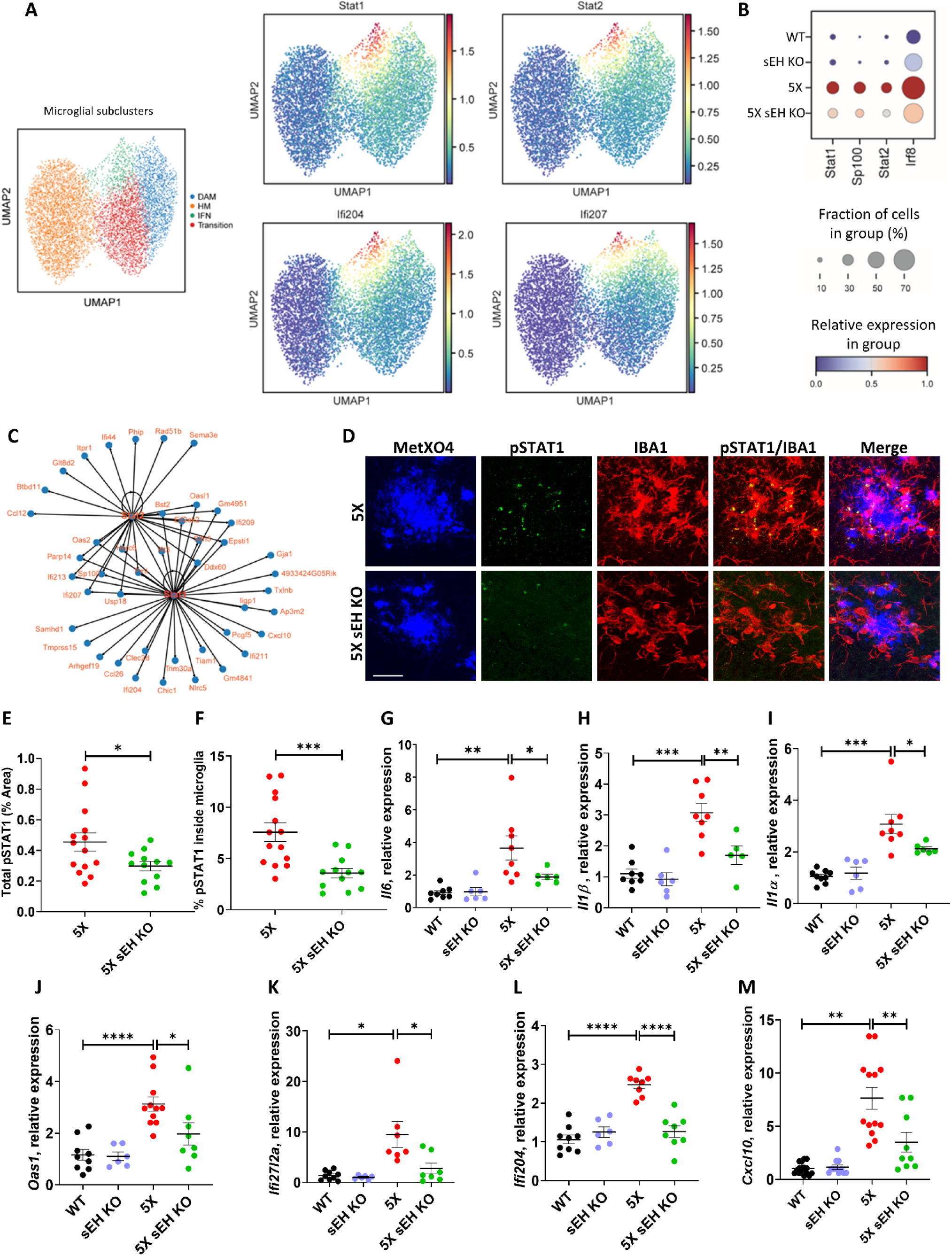
sEH deletion suppresses interferon-responsive gene networks and STAT1 activation. (A) UMAP of reclustered microglial nuclei identifying major microglial subclusters, including disease-associated microglia (DAM), interferon-responsive microglia (IFN), homeostatic microglia (HM), and transitional states. Feature plots show expression of interferon-associated transcription factors (*Stat1*, *Stat2*) and interferon-stimulated genes (*Ifi204*, *Ifi207*) overlaid on microglial subclusters. (B) Dot plot displaying the relative expression and fraction of cells expressing IFN-related transcription factors (*Stat1*, *Stat2*, *Sp100*, *Irf8*) across genotypes (WT, sEH KO, 5XFAD, and 5XFAD sEH KO). (C) SCENIC analysis of single-nucleus RNA-sequencing (snRNA-seq) data identified a microglial IFN-associated regulatory network centered on the transcription factors *Stat1* and *Stat2*, with connections to multiple IFN-responsive and immune-related genes. The network illustrates the inferred regulatory relationships between these transcription factors and their potential target genes. (D) Representative hippocampal immunofluorescence images from 5XFAD (5X) and 5X sEH KO mice stained for Methoxy X04 (β-amyloid, blue), phosphorylated STAT1 (pSTAT1, green), and IBA1 (microglia, red), with merged panels shown. (E–F) Quantification showing (E) total pSTAT1-positive area and (F) percentage of pSTAT1 signal colocalized with microglia. Each symbol represents analysis per image from a total of 14 brain sections from 3 mice in 5X and 12 sections in 5X sEH KO. (G–M) qPCR analysis of inflammatory and interferon-stimulated gene expression in hippocampal tissue, including *Il6* (G), *Il1b* (H), *Il1a* (I), *Oas1* (J), *Ifi27l2a* (K), *Ifi204* (L), and *Cxcl10* (M), across WT, sEH KO, 5XFAD, and 5XFAD sEH KO groups. Each point represents one mouse. Data are mean ± SEM. E&F: Welch’s *t*-test; G-M: unpaired t-tests with Welch’s correction were performed to compare WT vs 5X and 5X vs. 5X sEH KO to accommodate the unequal sample sizes and unequal variances present between groups. \**p* < 0.05, \*\**p* < 0.01, \*\*\**p* < 0.001, \*\*\*\**p* < 0.0001. Scale bar: 25 µm.

To further characterize the relationships among microglial subclusters, we performed pseudotime trajectory analysis, which positioned HM at one end of the trajectory with DAM and IFN microglia occupying more distal states (Fig. S3A). RNA velocity analysis supported directional transcriptional progression from HM toward IFN states (Fig. S3B). Differential gene expression analysis comparing microglia from 5XFAD versus WT mice identified robust induction of inflammatory and immune-associated genes (Fig. 1F). Gene ontology (GO) analysis of genes upregulated in 5XFAD microglia revealed significant enrichment of immune-related biological processes, including innate immune response, interferon signaling, and lysosomal pathways (Fig. 1G). Consistent with subcluster-specific analyses, DAM versus HM comparison showed enrichment of endolysosomal, phagocytic, and immune regulatory pathways (Fig. S3, C & D), whereas IFN versus HM comparison revealed strong enrichment for Type I interferon signaling and cytokine-related processes (Fig. S3, E & F). Differential expression analysis comparing 5XFAD sEH KO versus 5XFAD microglia demonstrated widespread downregulation of genes associated with inflammatory and immune activation (Fig. 1H). GO analysis of genes downregulated in 5XFAD sEH KO microglia revealed enrichment for pathways related to innate immune response, interferon signaling, and cytokine-mediated processes (Fig. 1I).

Collectively, these data indicate that germline disruption of sEH attenuates amyloid-associated remodeling of microglial transcriptional states, shifting microglia away from DAM and IFN-responsive programs.

### sEH disruption suppresses IFN gene networks and STAT1 activation

Given the reduction of IFN-responsive microglial populations in 5XFAD sEH KO mice, we next examined transcriptional programs associated with IFN activation. Visualization of canonical interferon-stimulated genes (ISGs), including *Ifi204* and *Ifi207*, demonstrated enrichment of these transcripts within the IFN-responsive microglial cluster (Fig. 2A).

To further define transcriptional regulators associated with this program, we examined IFN-related transcription factors. Dot-plot analysis revealed increased expression of *Stat1*, *Stat2*, *Irf8* and *Sp100* in microglia from 5XFAD mice compared with WT controls, which was markedly reduced in 5XFAD sEH KO mice (Fig. 2B). Consistent with this, regulatory network analysis identified STAT-centered gene modules encompassing multiple ISGs and immune-responsive genes that were prominently enriched in the 5XFAD condition and attenuated in the absence of sEH (Fig. 2C).

We next assessed whether these transcriptional changes were reflected at the protein level. Immunofluorescence staining for phosphorylated STAT1 (pSTAT1) revealed robust signal in IBA1⁺ microglia surrounding Methoxy-X04-positive plaques in 5XFAD mice, whereas pSTAT1 immunoreactivity was markedly reduced in plaque-associated microglia in 5XFAD sEH KO mice (Fig. 2D). Quantification confirmed a significant reduction in both total pSTAT1-positive area (Fig. 2E) and the percentage of pSTAT1 inside microglia (Fig. 2F) in 5XFAD sEH KO mice.

To determine whether suppression of IFN signaling was accompanied by reduced expression of downstream inflammatory mediators, we next measured cytokine and ISG expression. Quantitative PCR analysis demonstrated significant upregulation of pro-inflammatory cytokine genes, including *Il1α*, *Il1β* and *Il6*, in 5XFAD mice relative to WT controls, and these increases were attenuated in 5XFAD sEH KO mice (Fig. 2, G–I). Similarly, expression of ISGs, including *Ifi27l2a* and *Oas1*, and IFN-associated transcripts, including *Ifi204*, and *Cxcl10*, was significantly elevated in 5XFAD mice and attenuated following sEH disruption (Fig. 2, J-M).

These data demonstrate that sEH disruption attenuates the IFN-responsive microglial state, suppresses IFN-associated transcriptional networks and downstream STAT1 activation.

### 11,12-EET suppresses microglial STING activation and downstream interferon responses

Because the cGAS-STING pathway is a major driver of IFN responses, we assessed STING activation in vivo. Immunofluorescence analysis revealed prominent accumulation of phosphorylated STING (pSTING) in IBA1⁺ microglia within the hippocampus of 5XFAD mice, whereas both total pSTING-positive area and the percentage of pSTING inside microglia were reduced in sEH-deficient animals (Fig. 3, A-C).

**Figure 3.**
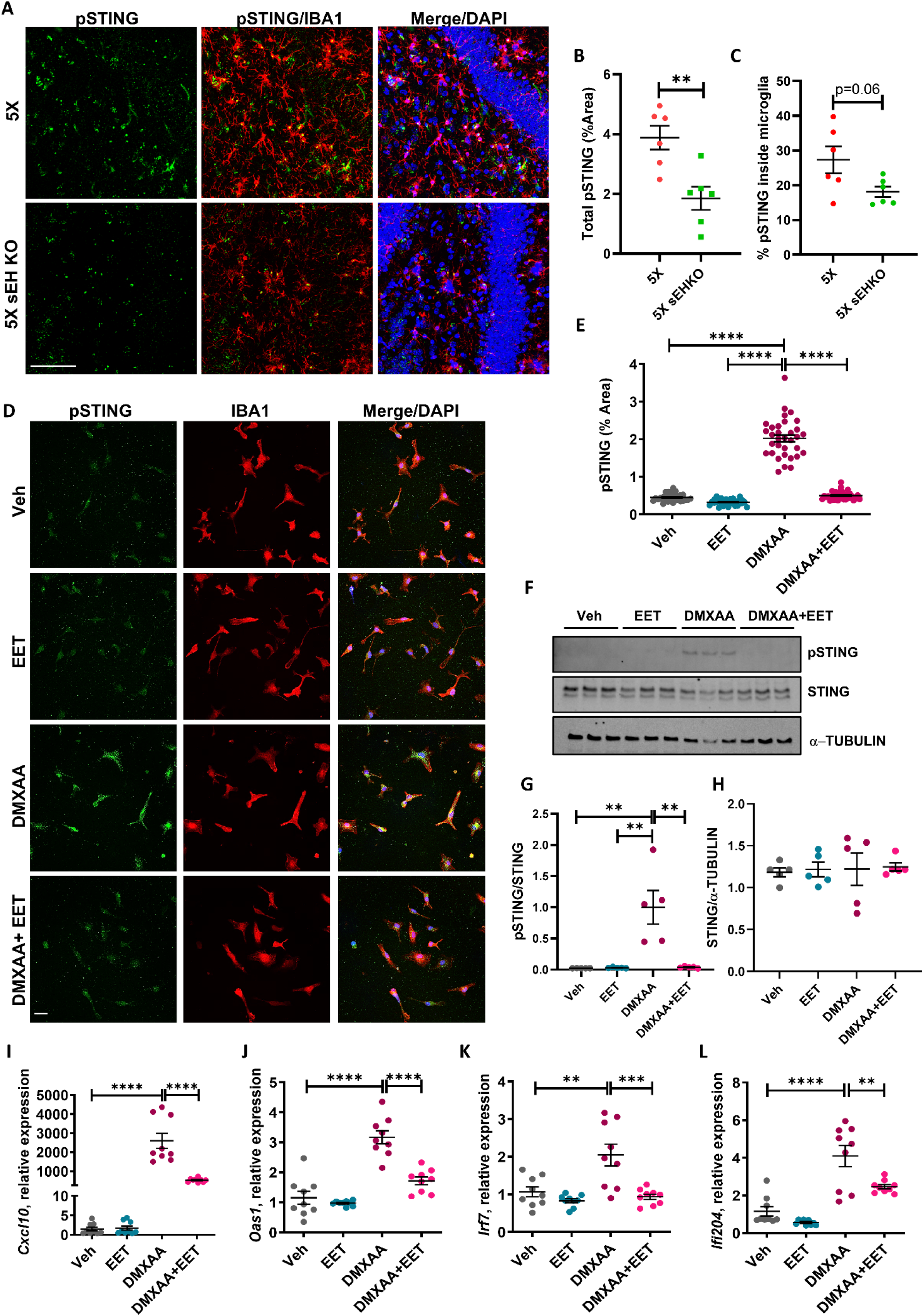
EET suppresses microglial STING activation and downstream interferon responses. (A) Representative hippocampal immunofluorescence images from 6-month-old 5XFAD (5X) and 5XFAD sEH knockout (5X sEH KO) mice stained for phosphorylated STING (pSTING, green), IBA1 (microglia, red), and DAPI (blue), with merged panels as indicated. (B-C) Quantification of total pSTING signal (% area) and percentage of pSTING inside microglia are shown in (B) and (C). Each point represents one mouse. (D-E) Representative immunofluorescence images of primary microglia treated with vehicle (Veh), EET, DMXAA (STING agonist), or DMXAA + EET, stained for pSTING (green), IBA1 (red), and DAPI (blue). (D). Quantification of pSTING intensity (% area) is shown in (E). Each point represents one field from independent cultures. (F-H) Western blot analysis of pSTING and total STING in primary microglia treated as indicated, with α-TUBULIN as a loading control (F). Quantification of the pSTING/STING (G) and STING/α-TUBULIN (H) ratio is shown. (I–J) qPCR analysis of interferon-stimulated genes (ISGs) including *Cxcl10* (I), *Oas1* (J), *Irf7* (K), and *Ifi204* (L) following treatment with vehicle, EET, DMXAA, or DMXAA + EET in primary microglia; data are mean ± SEM. Statistics: Welch’s *t*-test (B&C); One-way ANOVA followed by Tukey’s post hoc test (E, G-L). \**p* < 0.05, \*\**p* < 0.01, \*\*\**p* < 0.001. Scale bar: 25 µm.

Because sEH is primarily expressed in astrocytes and regulates the availability of EETs, we next examined whether EETs can modulate STING activation in microglia. Primary microglia were treated with vehicle, 11,12-EET (abbreviated as EET hereafter), STING agonist DMXAA, or DMXAA together with EET (Fig. 3D). Under basal conditions, pSTING levels remained low in vehicle- and EET-treated cultures. DMXAA induced robust pSTING accumulation, whereas co-treatment with EET markedly attenuated DMXAA-induced pSTING signal (Fig. 3, D & E).

Consistent with the immunofluorescence findings, immunoblot analysis demonstrated increased pSTING levels following DMXAA stimulation, which were significantly reduced by EET treatment, while total STING protein levels remained unchanged (Fig. 3, F-H). These findings indicate that EET suppresses STING phosphorylation in response to agonist stimulation.

Consistent with pSTING levels, DMXAA induced robust expression of canonical ISGs, including *Cxcl10*, *Oas1*, *Irf7*, and *Ifi204*, while co-treatment with EET significantly attenuated induction of all four ISGs (Fig. 3, I-L), Together, these findings demonstrate that EET suppresses STING activation and downstream interferon-responsive transcriptional programs in microglia.

### Lysosomal involvement in EET-mediated STING suppression

Since activated STING traffics to lysosomes for degradation and signal termination, we next examined whether EET-mediated suppression of STING is lysosome-dependent. DMXAA induced robust pSTING accumulation in primary microglia and promoted localization of pSTING within LAMP1-positive compartments (Fig. 4A). Blockade of lysosomal acidification with Bafilomycin A1 (BAF) further enhanced DMXAA-induced pSTING levels, consistent with impaired lysosomal degradation of activated STING, the effect of both were attenuated by EET treatment (Fig. 4, A and B). These findings indicate that lysosomes serve as a critical checkpoint for STING signal termination and is required for EET-mediated suppression of STING activation.

**Figure 4.**
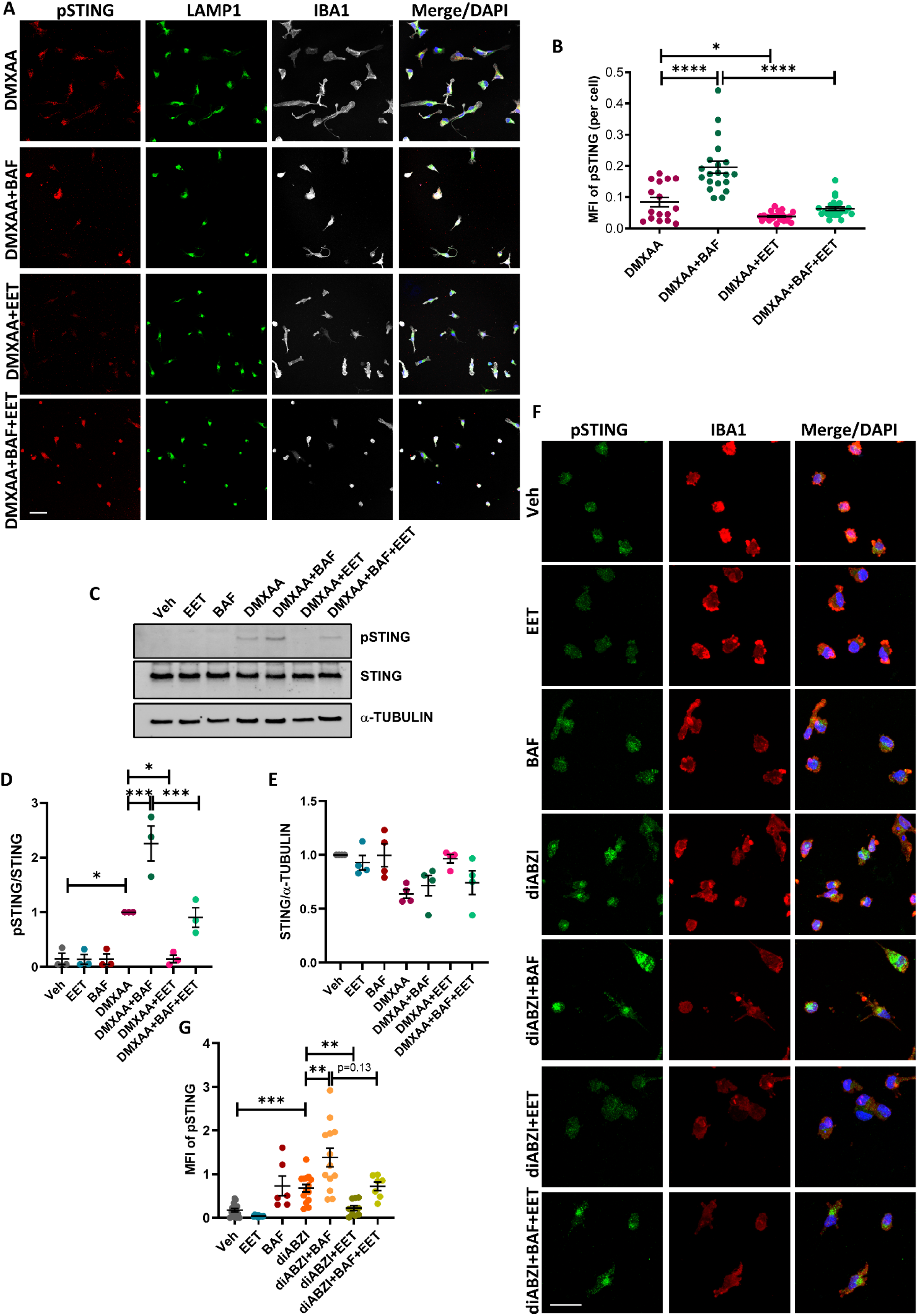
Lysosomal acidification is required for EET-mediated STING suppression. (A) Representative immunofluorescence images of primary microglia treated with DMXAA, DMXAA + bafilomycin A1 (BAF), DMXAA + EET, or DMXAA + EET + BAF and stained for phosphorylated STING (pSTING, red), LAMP1 (lysosomes, green), and IBA1 (microglia, gray), with DAPI nuclear counterstain (blue). (B) Quantification of pSTING mean fluorescence intensity (MFI) per cell. (C) Western blot analysis of pSTING and total STING in primary microglia treated with vehicle (Veh), EET, BAF, DMXAA, DMXAA + BAF, DMXAA + EET and DMXAA + BAF + EET, with α-TUBULIN as a loading control. (D, E) Quantification of the pSTING/STING ratio (D) and total STING/α-TUBULIN ratio (E). (F) Representative immunofluorescence images of human iPSC-derived microglia treated with vehicle (Veh), EET, BAF, diABZI, diABZI + BAF, diABZI + EET, or diABZI + BAF + EET and stained for pSTING (green), IBA1 (red), and DAPI (blue). (G) Quantification of pSTING mean fluorescence intensity per cell. Each point represents an individual field. Data are mean ± SEM. One-way ANOVA followed by Tukey’s post hoc test (B, D, E) & Welch’s ANOVA followed by Games-Howell test for G. \**p* < 0.05, \*\**p* < 0.01, \*\*\**p* < 0.001. Scale bar: 25 µm.

To independently validate these findings, STING activation was assessed by immunoblot analysis. Consistent with the immunofluorescence data, DMXAA treatment induced a marked increase in pSTING levels, whereas EET co-treatment substantially reduced STING phosphorylation (Fig. 4, C and D). Inhibition of lysosomal acidification with BAF increased pSTING accumulation following DMXAA stimulation, levels of which were reduced upon EET treatment (Fig. 4D). Total STING protein levels were not significantly altered across treatment groups (Fig. 4E), suggesting that EET primarily regulates STING activation and signal resolution rather than overall STING abundance. Together, these results demonstrate that lysosomal activity modulates EET-mediated attenuation of STING signaling.

We next investigated whether this mechanism is conserved in human microglia. Human iPSC-derived microglia were treated with vehicle, EET, BAF, the human-selective STING agonist diABZI, diABZI + EET, diABZI + BAF, or diABZI + EET + BAF. Similar to primary mouse microglia, diABZI induced robust pSTING accumulation that was markedly attenuated by EET treatment (Fig. 4, F & G). Inhibition of lysosomal acidification reversed pSTING accumulation in diABZI + EET-treated cells, demonstrating that lysosomal function is required for EET-dependent control of STING signaling across species.

### EET enhances microglial phagocytosis and lysosomal hydrolase activity

Given that EET suppresses STING activation and interferon-stimulated gene expression in microglia, we next examined whether EET treatment also influences microglial phagocytic function and lysosomal activity, two processes closely linked to amyloid clearance and inflammatory state regulation in neurodegenerative disease. Primary microglia were treated with vehicle or EET, and phagocytosis was evaluated using fluorescent bead uptake or pH-sensitive Aβ probes.

Immunofluorescence analysis revealed increased uptake of Alexa fluor (AF) 488-labeled beads in EET-treated microglia compared with vehicle-treated controls (Fig. 5, A & B). Consistent with these observations, treatment with pHrodo-conjugated Aβ42, which becomes fluorescent upon delivery to acidic compartments, revealed increased pHrodo-Aβ fluorescence intensity in EET-treated microglia 1 hour after pHrodo-Aβ incubation (Fig. 5, C & D). To determine whether enhanced Aβ uptake resulted in more efficient degradation, pHrodo-Aβ was measured 24 hours later, which showed marked reduction in EET treated cells (Fig. 5E). This result was further validated by assessing FAM labelled Aβ42 following 24 hours of incubation in vehicle or EET treated microglial cultures, showing significantly lower levels of intracellular Aβ42 by EET treatment (Fig. S4, A & B). Together, these data indicate that EET treatment enhances both phagocytic uptake and downstream processing of Aβ in primary microglia, in line with prior studies linking microglial activation state and lysosomal function to amyloid clearance efficiency [8, 9, 43].

**Figure 5.**
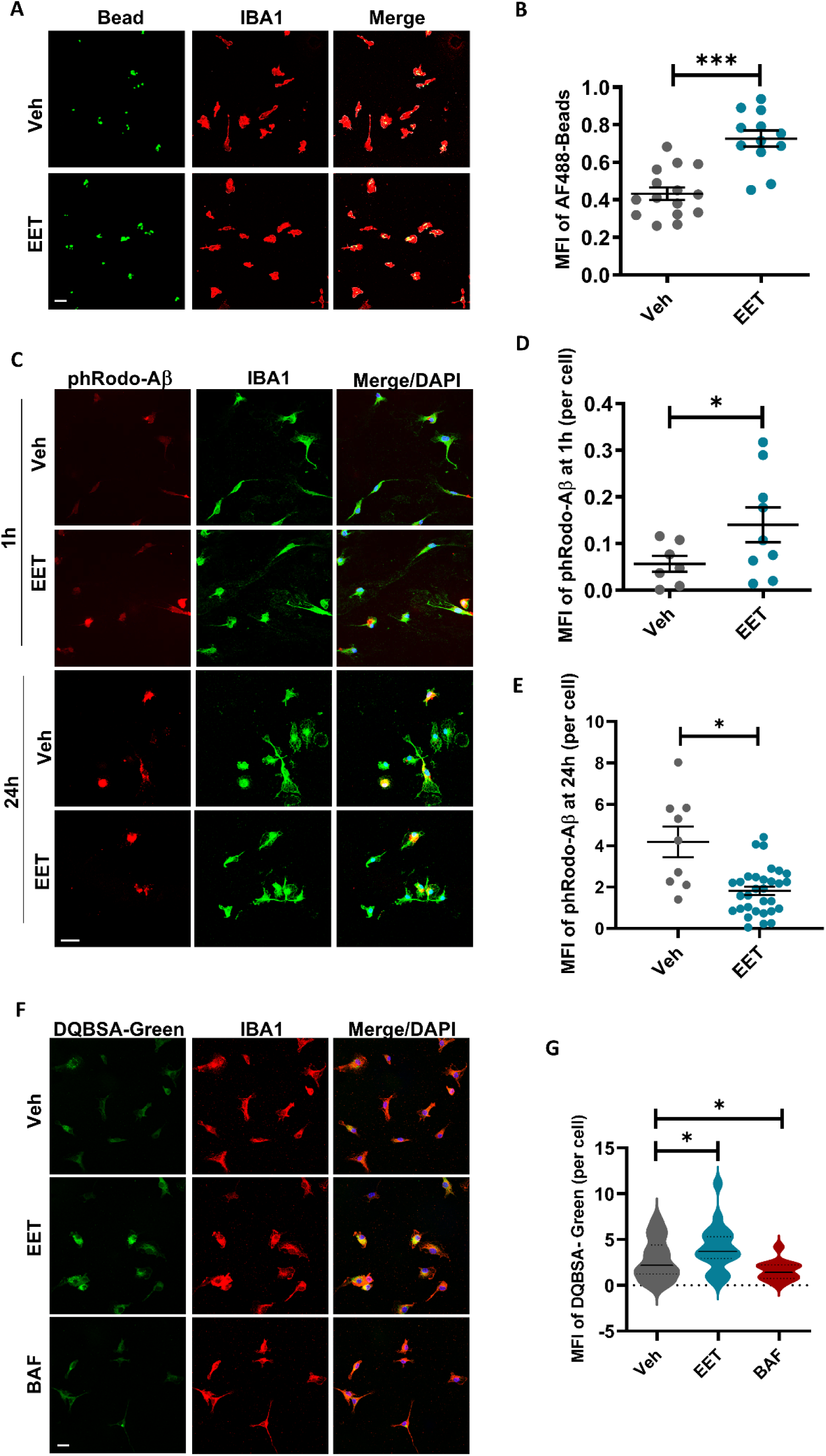
EET enhances microglial Aβ phagocytosis and promotes lysosomal activity. (A) Representative images of primary microglia treated with vehicle (Veh) or EET, showing uptake of fluorescent beads (green) in IBA1⁺ microglia (red), with merged panels as indicated. (B) Quantification of mean fluorescence intensity (MFI) of AF488 beads per cell. (C) Representative images of primary microglia treated with vehicle (Veh) or EET and incubated with pHrodo-red Aβ (red) for 1h, washed, fixed and imaged at indicated time points (1 h and 24 h), with IBA1 (green) and DAPI (blue) staining. (D, E) Quantification of pHrodo-Aβ fluorescence per cell at 1 h (D) and 24 h (E). (F) Representative images of primary microglia treated with vehicle (Veh), EET, or bafilomycin A1 (BAF), showing lysosomal proteolysis assessed by DQ-BSA Green (green) and IBA1 (red), with merged panels. (G) Quantification of DQ-BSA fluorescence per cell. Each point represents an individual field; data are mean ± SEM. (B, D, E): Welch’s *t*-test; G: unpaired *t*-tests with Welch’s correction were performed to compare Veh vs EET and Veh vs BAF to accommodate the unequal sample sizes and unequal variances present between groups \**p* < 0.05, \*\*\**p* < 0.001 where indicated. Scale bar: 25 µm.

We next assessed whether EET treatment modulates lysosomal hydrolase activity, a key determinant of cargo degradation capacity. Microglia were incubated with DQ-BSA Green, a self-quenched proteolytic substrate that fluoresces upon lysosomal degradation. EET treatment increased DQ-BSA fluorescence relative to vehicle, whereas inhibition of lysosomal acidification with bafilomycin A1 (BAF) markedly reduced the signal (Fig. 5, F and G). Collectively, these findings demonstrate that EET enhances microglial phagocytosis and lysosomal hydrolase activity, supporting a model in which improved lysosomal competence contributes to both enhanced Aβ clearance and suppression of STING activity and innate immune signaling.

### Astrocyte-specific disruption of sEH reduced amyloid pathology and STING signaling

Previous studies have shown that sEH is highly expressed in astrocytes, but can also be detected in brain endothelial cells where it regulates vascular function [36, 55, 56]. We next sought to determine the cellular source responsible for the protective effects observed in germline sEH-deficient mice by crossing the floxed *Ephx2* mice with GFAP-Cre or Cdh5-Cre to generate astrocytic (5X sEH AKO) or endothelial (5X sEH EKO) disruption of sEH on 5XFAD background [40–42].

Immunoblot analysis confirmed efficient deletion of sEH in 5XFAD astrocyte-specific knockout mice (5X sEH AKO) (Fig. 6, A & B). Methoxy-X04 staining revealed reduced amyloid plaque burden in AKO mice relative to 5XFAD controls (Fig. 6, C-F). Quantification demonstrated significant reductions in plaque number and size, accompanied by a trend toward reduced total plaque area. Additionally, to assess the association between microglia and Aβ plaques, we performed a spatial analysis of microglia residing within 20 µm of the plaque surface. Quantification revealed a significant reduction in the absolute count of plaque-proximal microglia in 5X sEH AKO compared with 5XFAD controls, indicating reduced microglial accumulation around amyloid deposits (Fig. 6G). Consistent with reduced pathology, D54D2 staining revealed decreased amyloid deposition together with reduced IBA1 and GFAP immunoreactivity in AKO mice (Fig. 6, H-K), indicating attenuation of plaque-associated gliosis. In contrast astrocytic sEH knockout, endothelial-specific disruption of sEH did not significantly alter plaque burden, amyloid deposition, microgliosis, or astrogliosis (Fig. S5).

**Figure 6.**
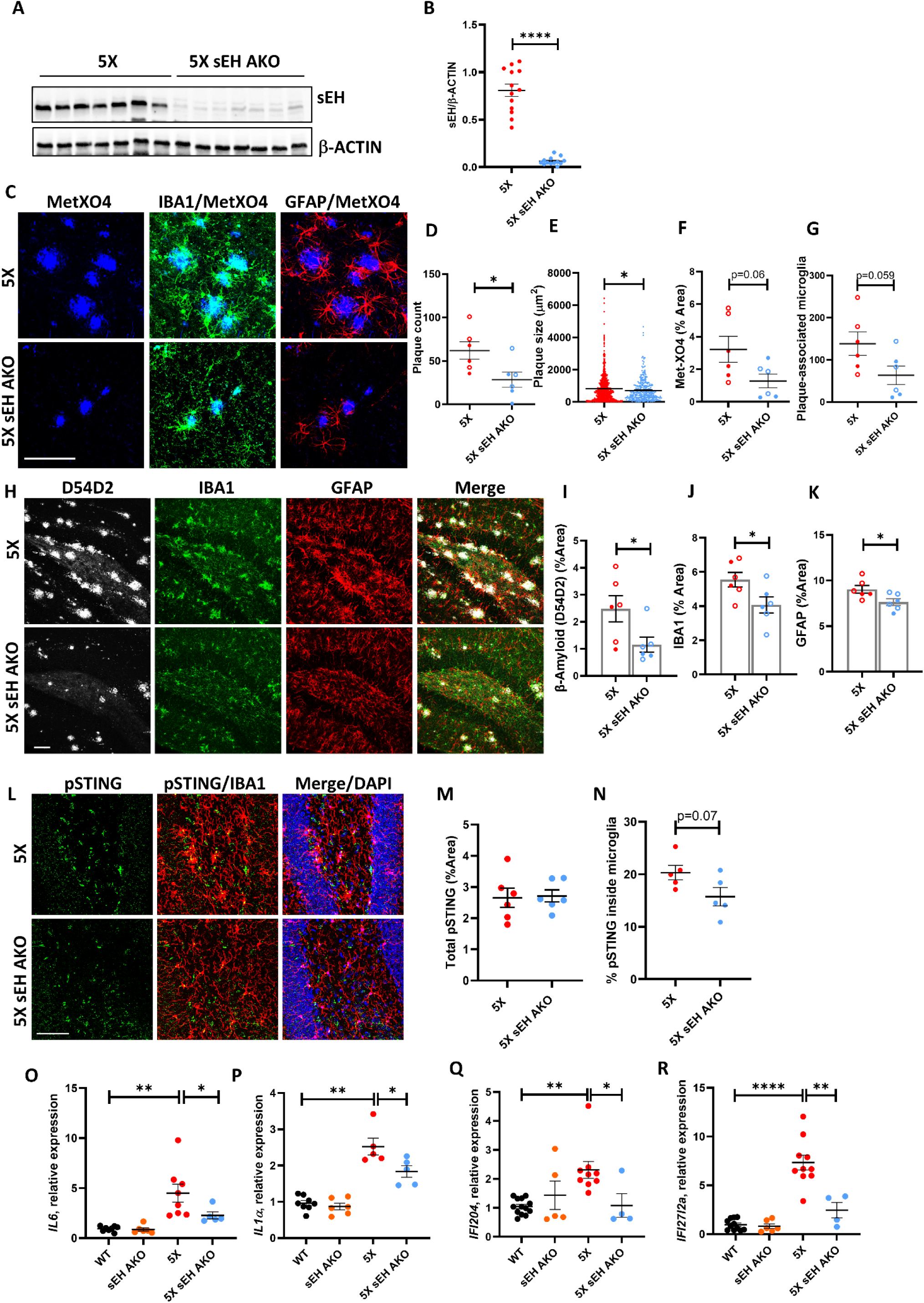
Astrocyte-specific deletion of sEH recapitulates reduced pathology and STING signaling. (A) Western blot analysis of sEH levels from hippocampal lysates of 5XFAD (5X) and astrocyte-specific sEH knockout (5X sEH AKO) mice, with β-ACTIN as a loading control. (B) Quantification of sEH/β-ACTIN ratio. (C) Representative confocal images of cortical sections stained with Methoxy X04 (MetX04; β-amyloid, blue). (D-G) Quantification of (D) absolute plaque counts, (E) plaque size, (F) MetX04-positive area, and (G) the number of plaque-associated microglia per image field. Plaque counts were determined by surface rendering in Imaris. For each mouse, measurements were averaged across three sections. (H) Representative confocal images of hippocampal sections stained with D54D2 (β-amyloid, white), IBA1 (microglia, green), and GFAP (astrocytes, red), with merged panels as indicated. (I-K) Quantification of D54D2-positive area (I), IBA1-positive area (J), and GFAP-positive area (K). (L) Representative hippocampal immunofluorescence images stained for phosphorylated STING (pSTING, green), IBA1 (microglia, red), and DAPI (blue), with merged panels as indicated. (M, N) Quantification of total pSTING-positive area (M) and percentage of pSTING-positive microglia (N). (O-R) qPCR analysis of inflammatory and interferon-responsive genes including *Il6* (O), *Il1α* (P), *Ifi204* (Q), and *Cxcl10* (R) in hippocampal tissue from WT, sEH AKO, 5XFAD, and 5X sEH AKO mice. Each symbol represents one mouse; data are mean ± SEM. Statistics: B, D-G, I-K, M, N, O-R: Welch’s *t*-test; \**p* < 0.05, \*\**p* < 0.01, \*\*\*\**p* < 0.0001. Scale bar: 75 µm.

Given the reduction in IFN-responsive transcriptional programs observed in germline sEH-deficient mice, we next examined STING activation in astrocyte-specific knockout animals. Immunofluorescence staining revealed abundant pSTING immunoreactivity within IBA1⁺ microglia in 5XFAD mice (Fig. 6L). Quantification showed that total pSTING immunofluorescence intensity in the hippocampus was not significantly altered in 5X sEH AKO mice relative to 5XFAD controls (Fig. 6M). In contrast, the percentage of pSTING-positive microglia trended lower in astrocyte-specific knockout mice compared with 5XFAD controls (Fig. 6N), indicating selective attenuation of microglial STING activation. Further supporting a functional role of astrocytic sEH, hippocampal expression of inflammatory and interferon-responsive genes, including *Il6*, *Il1α*, *Ifi204*, and *Cxcl10*, was significantly reduced in 5X sEH AKO mice (Fig. 6, O-R). Together, these findings demonstrate that astrocytic, but not endothelial, sEH is a major contributor to amyloid pathology, glial activation, and microglial innate immune signaling in the 5XFAD brain, establishing an astrocyte-microglia axis through which the sEH-EET pathway regulates lysosomal and immune homeostasis during AD progression.

## Discussion

In this study, we extend our prior work revealing that genetic or pharmacologic sEH inhibition restores glial homeostasis in both amyloid and tau mouse models [36, 37]. We demonstrate here that sEH disruption in 5XFAD mice suppresses disease-associated and IFN-responsive microglial programs and restrained STAT1 activation. We provide mechanistic depth by identifying lysosome-dependent termination of STING signaling as the downstream effector that couples epoxy-lipid signaling to lysosomal competence and innate immunity in the AD brain.

Type I IFN signaling has emerged as a pathogenic amplifier in AD, linking amyloid accumulation to synaptic dysfunction and cognitive decline [14, 15, 57], and aberrant cGAS–STING activation drives neuroinflammation in aging and amyloidosis [17–19]. That sEH loss broadly repressed IFN-associated transcription factors and ISG networks, with reduced pSTING and pSTAT1 in plaque-associated microglia, places the sEH–EET pathway upstream of this cascade.

We show that EET treatment enhanced lysosomal hydrolase activity and degradation of phagocytosed cargo, whereas disruption of lysosomal acidification suppressed STING activation. Because lysosomes govern degradation of amyloid cargo while simultaneously serving as critical sites for STING signal termination, our data support a model in which lysosomal competence coordinately regulates inflammatory state and amyloid clearance. These findings align with previous observations that lysosomal dysfunction limits microglial clearance efficiency in AD and that restoration of lysosomal activity improves amyloid degradation [58–60]. Together, our data suggest that rather than representing independent processes, suppression of STING signaling and enhancement of Aβ degradation may arise from a common upstream mechanism involving restoration of lysosomal function by the sEH-EET axis.

Cell-type-specific deletion refines the cellular origin of this mechanism. Previous studies established protective effects of global sEH inhibition, but the responsible compartment remained unclear. Astrocytes are increasingly recognized as regulators of inflammatory tone, lipid homeostasis, and the extracellular milieu in neurodegeneration [61–63]. Although not considered primary phagocytes of amyloid, their capacity to shape lipid availability plausibly influences microglial responses at the tissue level. Consistent with this, astrocyte-specific disruption of sEH recapitulated many protective phenotypes of germline knockout mice, including reduced amyloid pathology, gliosis, and inflammatory gene expression, whereas endothelial-specific disruption had minimal effect. Total hippocampal pSTING was not significantly altered, though astrocyte-specific deletion showed a trend toward reduced microglial pSTING. Together these findings identify astrocytic sEH as a major regulator of brain EET availability and support an astrocyte-microglia axis through which epoxy lipid metabolism shapes microglial lysosomal competence, immune state, and amyloid clearance.

Recently, Wu et al. identified a liver-brain axis in which hepatic sEH regulates circulating EET levels and influences amyloid pathology, tauopathy, and cognition [39]. Together with our astrocyte-specific results, this suggests that central and peripheral pools of sEH contribute to disease progression through distinct but convergent EET-dependent pathways.

## Conclusions

Our findings support a model in which astrocyte sEH-EET axis shapes microglial responses to amyloid pathology through regulation of STING-IFN-STAT1 signaling. By placing the sEH-EET pathway upstream of both STING-dependent inflammatory signaling and Aβ degradation, this study suggests that lysosomal competence may coordinate immune state and proteostatic function in the diseased brain. These observations provide a framework for understanding how glial metabolic pathways influence neurodegeneration and support further investigation of sEH inhibition as a therapeutic strategy in Alzheimer’s disease.

## Acknowledgements

We thank Brittany Reeves and Haiying Liu for expert technical assistance, Yanyan Qi and Hongjie Li for snRNA-seq experiment, Xinlong Luo for meta data submission, and members of the Zheng lab for valuable comments and suggestions. We are grateful to Dr. Artiom Gruzdev of the NIEHS GEMM core facility for making the *Ephx2* floxed mice.

## Author Contributions

BG and HZ designed the overall study; BG performed major biological experiments with support from CSR on iPSC cultures, SW for immuno staining; CQ assisted with the snRNA-seq analysis; AG and MC performed initial analysis on germline sEH knockout mice; DCZ developed and provided the *Ephx2* floxed mice. All authors provided input, read, and approved the manuscript.

## Availability of data and materials

Raw snRNA-seq data have been deposited at NCBI GEO with the accession number: GSE351037 and are publicly available as of the date of publication. No original codes were generated in this study. Any additional materials or datasets used in the current study are available from the corresponding author on reasonable request.

## Declarations

### Ethics approval and consent to participate

All animal procedures were performed in accordance with NIH guidelines and with the approval of the Baylor College of Medicine Institutional Animal Care and Use Committee prior to all animal-related studies (IACUC protocol # AN-1853).

### Consent for publication

All authors have approved the contents of this manuscript and provided consent for publication.

## List of abbreviations

Aβ: Amyloid-beta
AD: Alzheimer’s disease
AF488: Alexa Fluor 488
AKO: Astrocyte-specific knockout
BAF: Bafilomycin A1
BBB: Blood-brain barrier
CD200: Cluster of Differentiation 200
cGAS: Cyclic GMP-AMP synthase
CX3CL1: C-X3-C motif chemokine ligand 1 (Fractalkine)
DAM: Disease-associated microglia
DHET: Dihydroxyeicosatrienoic acid
DMXAA: 5,6-Dimethylxanthenone-4-acetic acid
DQ-BSA: Dye-quenched bovine serum albumin
EET: Epoxyeicosatrienoic acid
EKO: Endothelial-specific knockout
ESCRT: Endosomal Sorting Complex Required for Transport
GFAP: Glial fibrillary acidic protein
GO: Gene Ontology
HPC: Hematopoietic progenitor cell
IBA1: Ionized calcium-binding adaptor molecule 1
IFN: Interferon
iMGL: Induced pluripotent stem cell-derived microglia-like cell
iPSC: Induced pluripotent stem cell
ISG: Interferon-stimulated gene
LAMP1: Lysosome-associated membrane protein 1
M-CSF: Macrophage colony-stimulating factor
MetX04: Methoxy-X04
OPC: Oligodendrocyte precursor cell
pSTAT1: Phosphorylated signal transducer and activator of transcription 1
pSTING: Phosphorylated stimulator of interferon genes
qPCR: Quantitative polymerase chain reaction
RNA-seq: RNA sequencing
SCENIC: Single-Cell Regulatory Network Inference and Clustering
sEH: Soluble epoxide hydrolase
snRNA-seq: Single-nucleus RNA sequencing
STAT1: Signal transducer and activator of transcription 1
STING: Stimulator of interferon genes
TGF-β1: Transforming growth factor-beta 1
TPPU: 1-Trifluoromethoxyphenyl-3-(1-propionylpiperidin-4-yl) urea
UMAP: Uniform manifold approximation and projection
WT: Wild type
5XFAD (5X): Five familial Alzheimer’s disease transgenic mouse model

