## Supplementary Methods, Figures, and Tables for "Astrocytic sEH–EET signaling regulates microglial lysosomal function and suppresses the STING–Type I interferon pathway in Alzheimer’s disease"

#### **Supplementary Methods**

##### **Behavioral analysis**

Behavioral testing was conducted with the experimenter blinded to genotype. Prior to each behavioral assay, mice were habituated to the testing room (150 lx ambient lighting, 60 dB white noise) for at least 30 min. A minimum of 24 h was allowed between different behavioral assays to permit recovery.

##### **Novel Object Recognition**

Novel object recognition (NOR) was performed in a Plexiglas arena (22 cm × 44 cm). During the training phase, mice were allowed to freely explore the arena containing two identical LEGO objects for 10 min. Twenty-four hours later, mice were returned to the same arena for the test phase, which consisted of one familiar object from the training phase and one novel object that differed in color and shape but was matched for size and volume. Mice were allowed to explore the objects freely for 10 min. Object exploration was defined as the mouse nose being $\leq 2$ cm from the object and was quantified using ANY-maze software. Recognition memory performance was expressed as the percentage of total exploration time spent investigating the novel object.

##### **Fear Conditioning**

The fear conditioning protocol consisted of a training phase followed by contextual and cued tests. During training, mice were placed in the conditioning chamber and allowed to explore freely for 2 min, after which an auditory conditioned stimulus (80 dB, 5 kHz tone) was presented for 30 s, immediately followed by a 2-s foot shock (0.8 mA). This pairing was repeated once, after which animals were returned to their home cages. The contextual fear test was conducted 24 h later by returning mice to the original training chamber for 5 min in the absence of both tone and shock. One hour later, the cued fear test was performed in a novel chamber with distinct geometric shape, flooring, lighting, and odor. After 3 min of exploration, the auditory stimulus was presented for an additional 3 min. Freezing behavior during all phases was quantified using FreezeFrame4 software (Actimetrics, Coulbourn Instruments) and expressed as the percentage of time spent freezing.

### Supplementary Figures

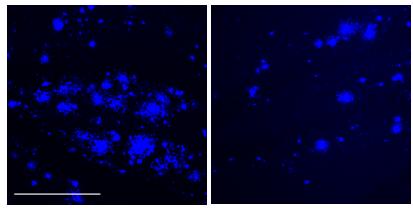

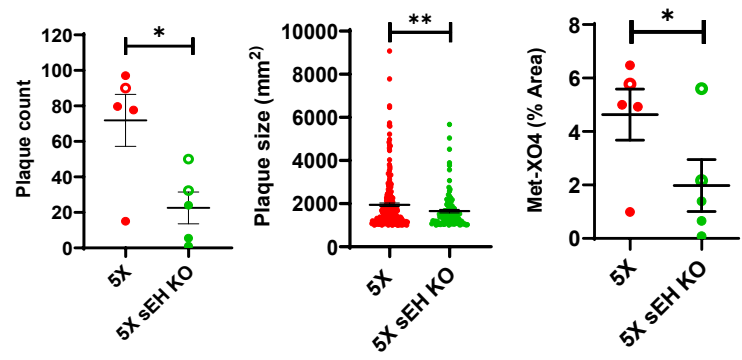

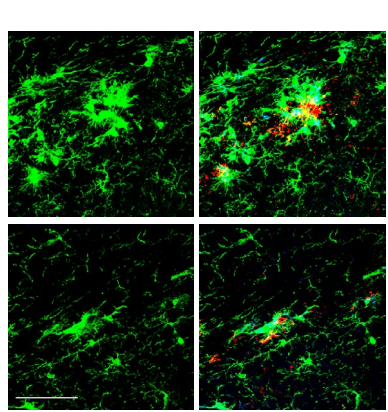

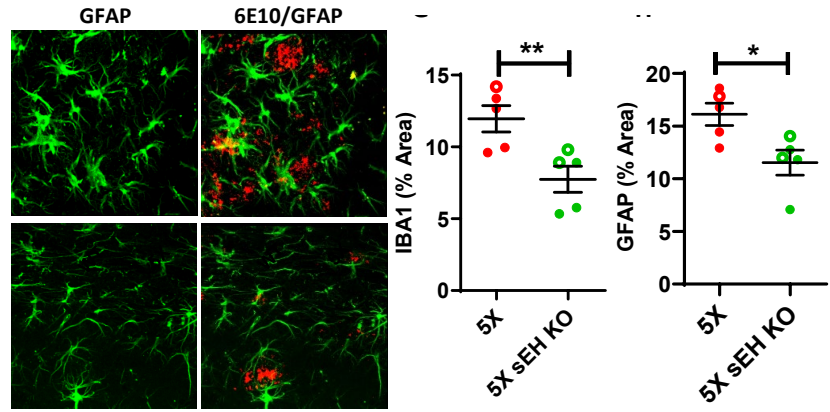

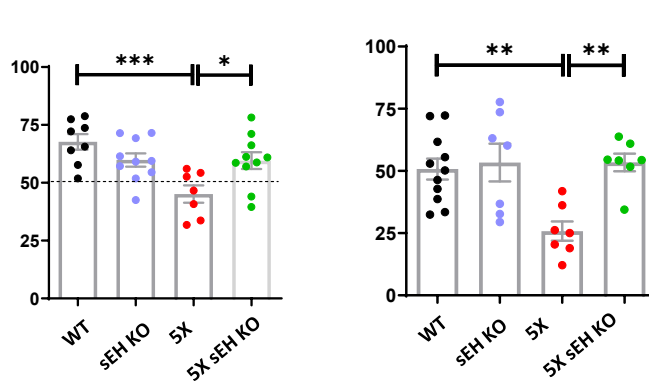

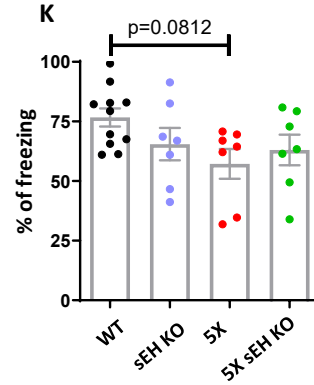

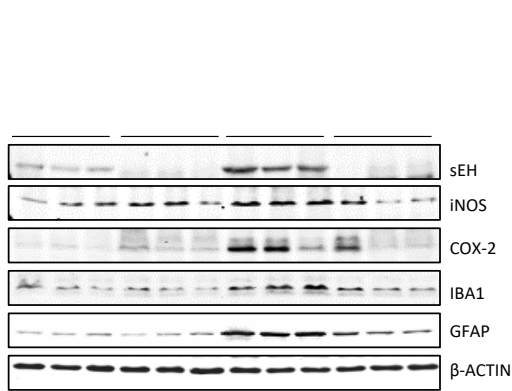

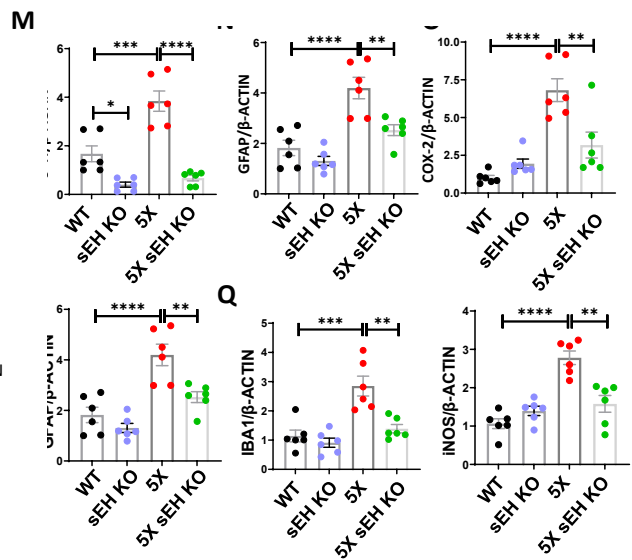

**Supplementary Figure 1. sEH deletion reduces amyloid plaque burden, gliosis, and improves cognitive function, and reduces inflammatory cytokines in 5XFAD mice.**

(A-D) Representative confocal images of hippocampal sections from 6-month-old 5XFAD (5X) and global 5XFAD sEH knockout (5X sEH KO) mice stained with Methoxy X04 (MetX04; β-amyloid, blue) (A). Quantification shows (B) plaque count, (C) plaque size (area, $\mu\text{m}^2$), and (D) MetX04-positive area (% area). (E-H) Representative images of plaque-associated microglia and astrocytes from 5X and 5X sEH KO mice stained for IBA1 (microglia, green), GFAP (astrocytes, green), and 6E10 (β-amyloid, red), with merged panels as indicated (E-F). Quantification shows (G) IBA1-positive area and (H) GFAP-positive area. (I-K) Behavioral assessment across WT, sEH KO, 5X, and 5X sEH KO mice. (I) Object discrimination index in the novel object recognition test. (J-K) Percentage freezing during contextual (J) and post-cue (K) fear conditioning. (L) Western blot analysis of hippocampal lysates showing expression of sEH, iNOS, COX-2, IBA1, and GFAP across WT, sEH KO, 5XFAD, and 5XFAD sEH KO mice, with β-ACTIN as a loading control. (M-R) Quantification of western blot analyses from panel (L), including sEH/β-ACTIN, COX-2/β-ACTIN, GFAP/β-ACTIN, IBA1/β-ACTIN, and iNOS/β-ACTIN ratios. Each symbol represents one mouse; data are mean ± SEM. Statistics: unpaired Welch's *t* test (B-D, G, H); one-way ANOVA with Tukey's multiple comparison test (I-K, M-R). \**p* < 0.05, \*\**p* < 0.01. Scale bar: 75 $\mu\text{m}$.

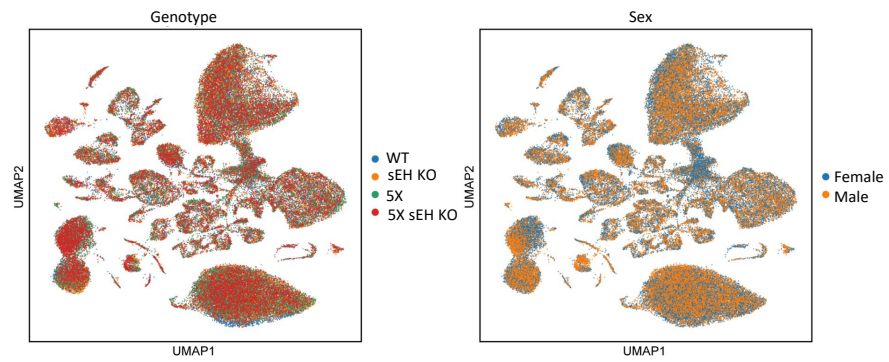

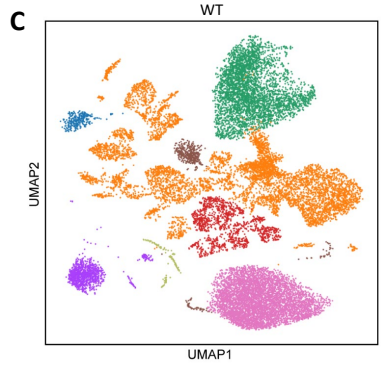

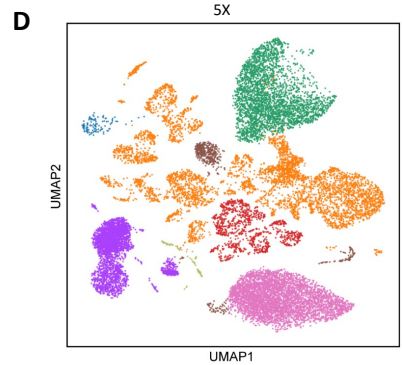

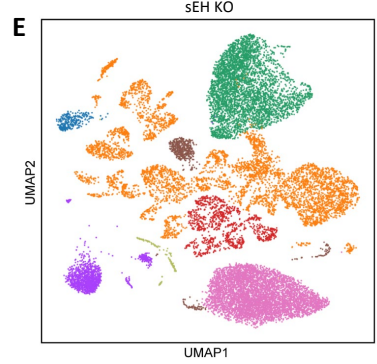

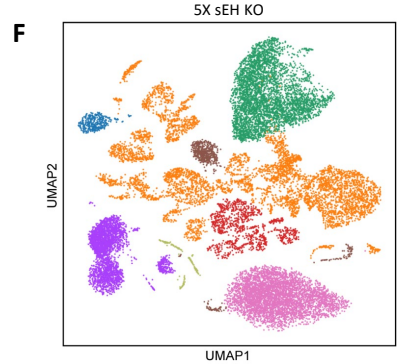

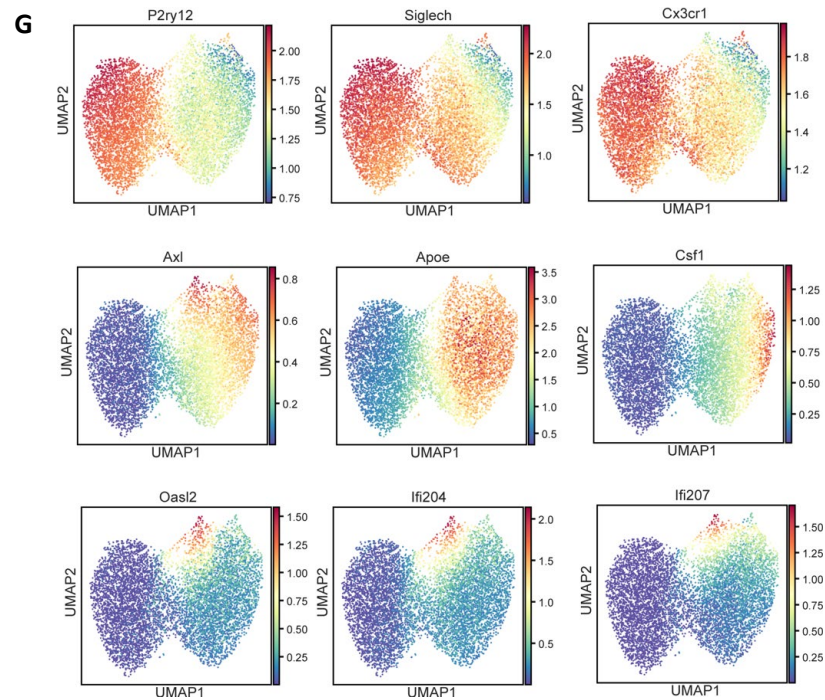

**Supplementary Figure 2. Batch correction, genotype distribution, and marker gene expression across snRNA-seq datasets.**

(A–B) UMAP plots of integrated single-nucleus RNA-seq datasets following batch correction, colored by genotype (A; WT, sEH KO, 5XFAD, 5XFAD sEH KO) and by sex (B; female and male). (C–F) UMAP plots of single-nucleus datasets separated by genotype, including WT (C), 5XFAD (D), sEH KO (E), and 5XFAD sEH KO (F), showing distribution of major cell types across groups. (G) Feature plots showing expression levels of representative marker genes across clusters, including *P2ry12*, *Siglech*, *Cx3cr1*, *Axl*, *Apoe*, *Csf1*, *Oas2*, *Ifi204*, and *Ifi207*, displayed as scaled expression intensity.

**A** Pseudotime Trajectory

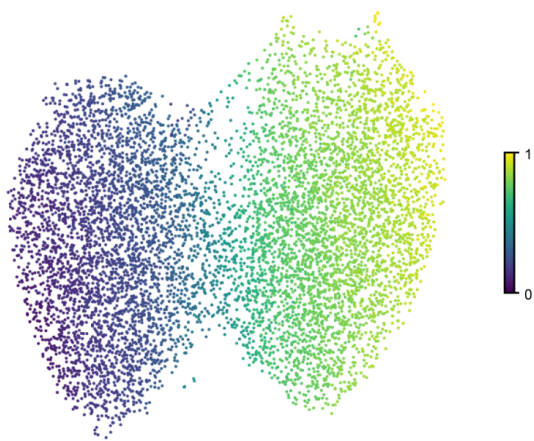

**B** RNA velocity

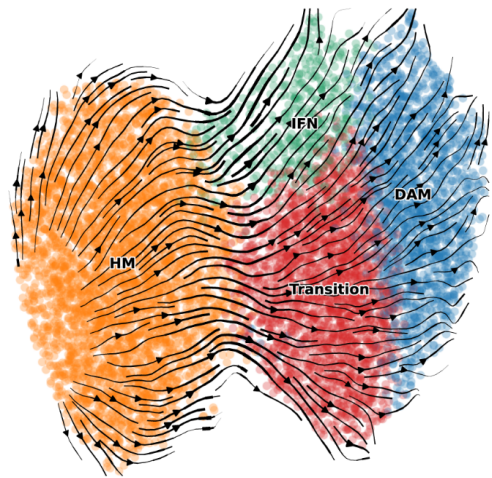

**C** DAM vs HM DEGs (microglial cells)

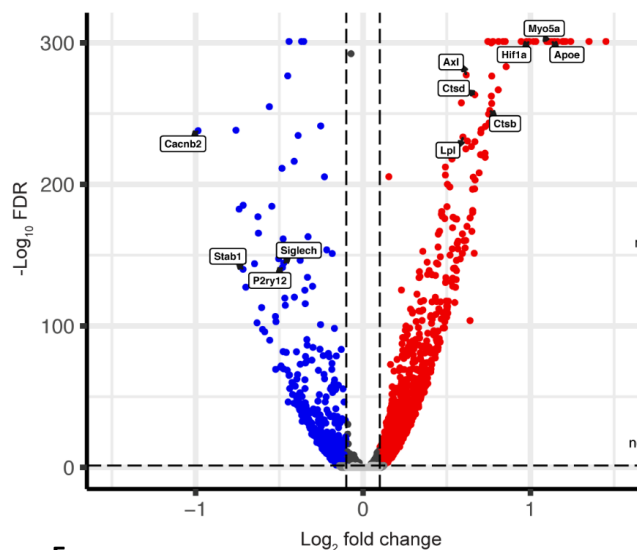

**D** GO enrichment for upregulated genes (DAM vs HM)

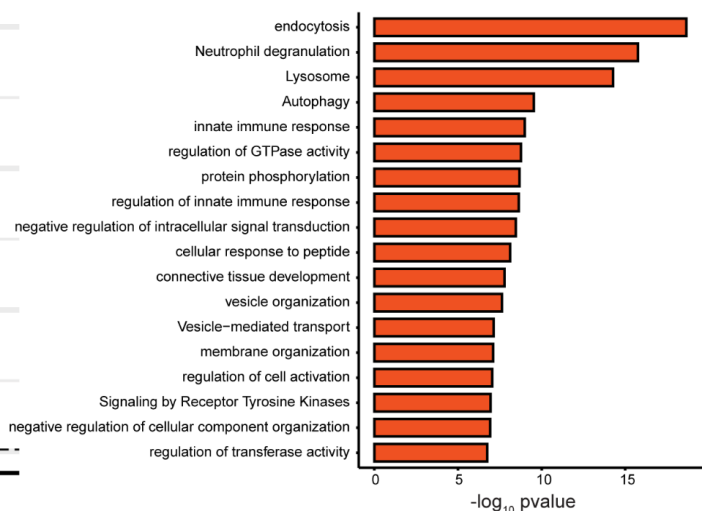

**E**

IFN vs HM DEGs (microglial cells)

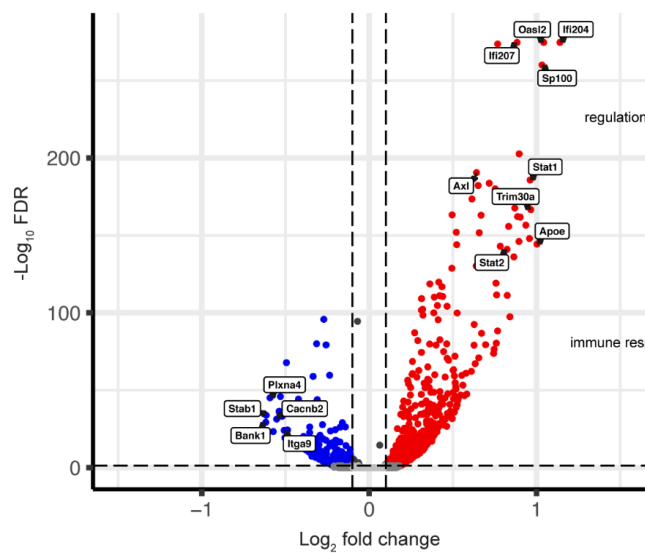

**F**

GO enrichment for upregulated genes (IFN vs HM)

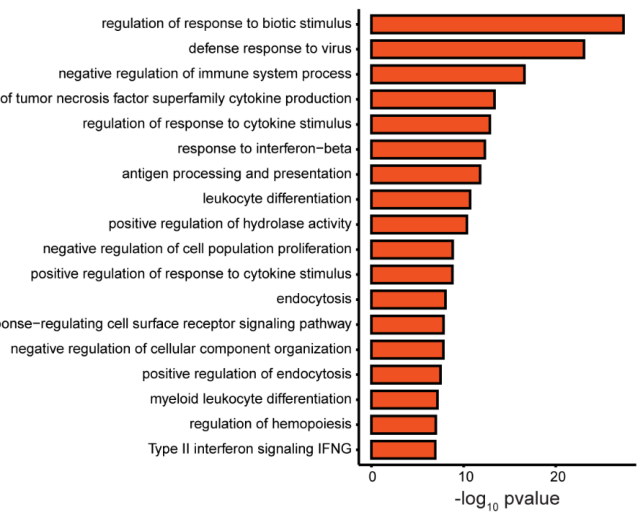

**Supplementary Figure 3. Pseudotime trajectory, RNA velocity, and differential gene expression analyses of microglial subclusters.**

(A) Pseudotime trajectory plot of microglial cells showing the inferred progression of transcriptional states across clusters, displayed as a continuous gradient. (B) RNA velocity projection overlaid on the UMAP embedding illustrating directional transcriptional dynamics across microglial subclusters, including homeostatic microglia (HM), disease-associated microglia (DAM), interferon-responsive microglia (IFN), and transitional states. (C) Volcano plot showing differentially expressed genes (DEGs) between DAM and HM microglia, with selected genes highlighted. (D) Gene Ontology (GO) enrichment analysis for genes upregulated in DAM compared with HM microglia, presented as $-\log_{10} p$ values for the top enriched biological processes. (E) Volcano plot showing differentially expressed genes between IFN and HM microglia, with representative genes indicated. (F) Gene Ontology (GO) enrichment analysis for genes upregulated in IFN microglia compared with HM microglia, showing the top enriched biological processes ranked by $-\log_{10} p$ value.

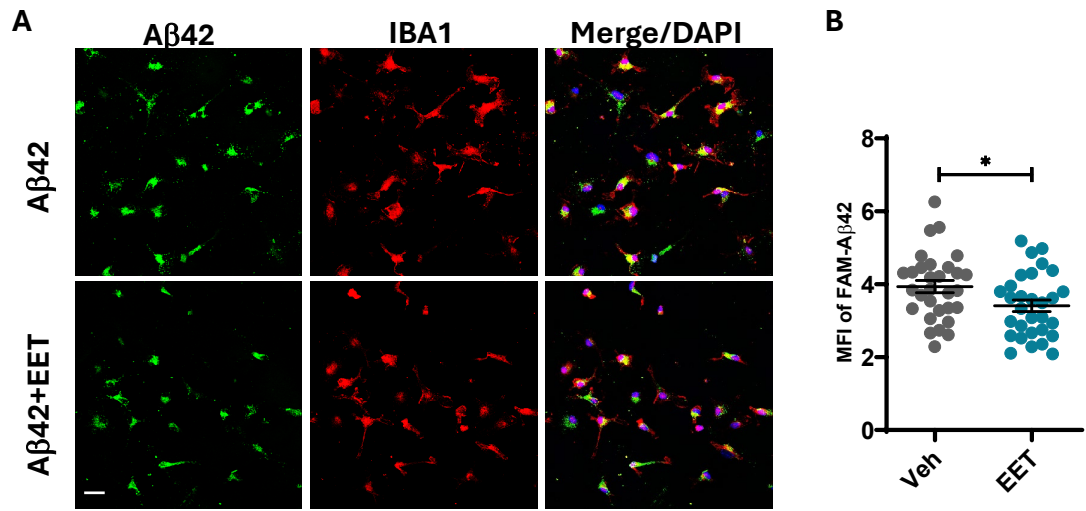

**Supplementary Figure 4. EET enhances Aβ clearance in primary microglia.**

(A) Representative immunofluorescence images of primary microglia incubated with FAM-Aβ42 (green) in the presence or absence of EET treatment, stained for IBA1 (microglia, red) and DAPI (blue), with merged panels as indicated. Insets show individual Aβ42 and IBA1 channels. (B) Quantification of Aβ42 mean fluorescence intensity (MFI) per cell in vehicle (Veh) and EET-treated microglia. Each point represents an individual field; data are mean ± SEM; unpaired Welch's *t* test. \*$p < 0.05$. Scale bar: 25 μm.

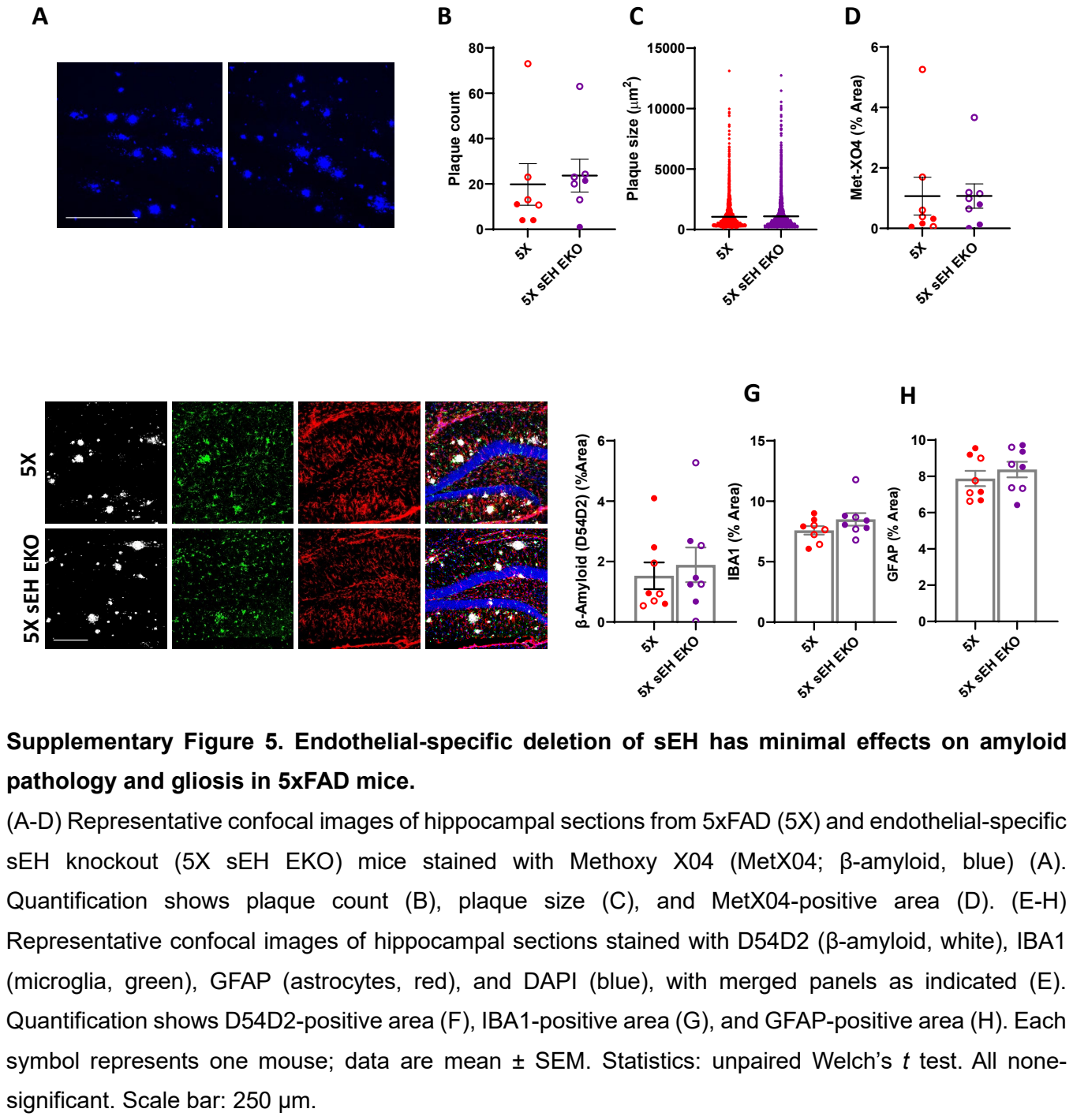

Supplementary Figure 5. Endothelial-specific deletion of sEH has minimal effects on amyloid pathology and gliosis in 5xFAD mice.

(A-D) Representative confocal images of hippocampal sections from 5xFAD (5X) and endothelial-specific sEH knockout (5X sEH EKO) mice stained with Methoxy X04 (MetX04; β-amyloid, blue) (A). Quantification shows plaque count (B), plaque size (C), and MetX04-positive area (D). (E-H) Representative confocal images of hippocampal sections stained with D54D2 (β-amyloid, white), IBA1 (microglia, green), GFAP (astrocytes, red), and DAPI (blue), with merged panels as indicated (E). Quantification shows D54D2-positive area (F), IBA1-positive area (G), and GFAP-positive area (H). Each symbol represents one mouse; data are mean ± SEM. Statistics: unpaired Welch’s t test. All none-significant. Scale bar: 250 µm.

**Supplementary Table S1. Primary antibodies used in this study**

| **Antibody** | **Host** | **Source** | **Identifier** | **Dilution** |
| --- | --- | --- | --- | --- |
| IBA1 | Rabbit | FUJIFILM<br>Wako | Cat# 01919741 | 1:500(IF)<br>1:1200(WB) |
| IBA1 | Goat | Novus<br>Biologicals | Cat# NB100-1028 | 1:500(IF) |
| GFAP | Mouse | Sigma-Aldrich | Cat# MAB360 | 1:500(IF)<br>1:1000(WB) |
| LAMP1 | Rat | BD<br>Pharmingen | Cat# 553792 | 1:500(IF) |
| pSTING(Ser365) | Rabbit | Cell<br>Signaling | Cat# 72971 | 1:100(IF)<br>1:1000(WB) |
| TMEM173/STING | Rabbit | Proteintech | 19851-1-AP | 1:10000(WB) |
| pSTING(Ser366) | Rabbit | Cell<br>Signaling | Cat# 50907 | 1:100(IF) |
| Aβ (D54D2) | Rabbit | Cell<br>Signaling | Cat# 8243 | 1:500(IF) |
| pSTAT1 (Tyr701) | Rabbit | Cell<br>Signaling | Cat# 9167 | 1:200(IF) |
| sEH | Rabbit | Bruce<br>Hammock |  | 1:1000(WB) |
| iNOS | Rabbit | Cell<br>Signaling<br>Technology | Cat# 2982 | 1:1000 |
| COX2 | Rabbit | Thermo<br>Fisher | Cat# 35-8200 | 1:500 |
| β-ACTIN | Mouse | Sigma-Aldrich | Cat#A5441 | 1:5000(WB) |
| α-TUBULIN | Mouse | Sigma-Aldrich | Cat# T6199 | 1:5000(WB) |
| GAPDH | Mouse | Invitrogen | Cat# AM4300 | 1:5000(WB) |

**Supplementary Table S2. Primers used in this study**

| **Gene** | **Forward (5'-3')** | **Reverse (3'-5')** |
| --- | --- | --- |
| Ifi204 | CCAGTCACCAATACTCCACAGC | CTCTGAGTGGAGAACAGCACCT |
| Cxcl10 | CCCACGTGTTGAGATCATTG | CACTGGGTAAAGGGGAGTGA |
| Irf7 | CCTCTGCTTTCTAGTGATGCCG | CGTAAACACGGTCTTGCTCCTG |
| Oas1 | GCTGCCAGCCTTTGATGT | TGGCATAGATTCTGGGATCA |
| Ifi27l2a | CTTCACTGGGACAGGCATTGCA | CCTGCTGATTGGAGTGTGGCTA |
| Il6 | TAGTCCTTCCTACCCCAATTTCC | TTGGTCCTTAGCCACTCCTTC |
| Il1a | CGCTTGAGTCGGCAAAGAAAT | CTTCCCGTTGCTTGACGTTG |
| Il1b | GCAACTGTTCCTGAACTCAACT | ATCTTTTGGGGTCCGTCAACT |
| Gapdh | AACAGCAACTCCCACTCTTC | CCTGTTGCTGTAGCCGTATT |
